# Antigenic mapping and functional characterization of human New World hantavirus neutralizing antibodies

**DOI:** 10.1101/2022.07.19.500579

**Authors:** Taylor B. Engdahl, Elad Binshtein, Rebecca L. Brocato, Natalia A. Kuzmina, Lucia M Principe, Steven A. Kwilas, Robert K. Kim, Nathaniel S. Chapman, Monique S. Porter, Pablo Guardado-Calvo, Felix A. Rey, Laura S. Handal, Summer M. Diaz, Irene A. Zagol-Ikapitte, Joseph X. Reidy, Andrew Trivette, Alexander Bukreyev, Jay W. Hooper, James. E. Crowe

## Abstract

Hantaviruses are high-priority emerging pathogens carried by rodents and transmitted to humans by aerosolized excreta or, in rare cases, person-to-person contact. While sporadic in North and South America, many infections occur in Europe and Asia, with mortality ranging from 1 to 40% depending on the hantavirus species. There are currently no FDA-approved vaccines or therapeutics for hantaviruses, and the only treatment for infection is supportive care for respiratory or kidney failure. Additionally, the humoral immune response to hantavirus infection is incompletely understood, especially the location of major antigenic sites on the viral glycoproteins and conserved neutralizing epitopes. Here, we report antigenic mapping and functional characterization for four neutralizing hantavirus antibodies. The broadly neutralizing antibody SNV-53 targets an interface between Gn/Gc, neutralizes through fusion inhibition and cross-protects against the Old World hantavirus species Hantaan virus when administered pre- or post-exposure. Another broad antibody, SNV-24, also neutralizes through fusion inhibition but targets domain I of Gc and demonstrates weak neutralizing activity across hantavirus species. ANDV-specific, neutralizing antibodies (ANDV-5 and ANDV-34) neutralize through attachment blocking and protect against hantavirus cardiopulmonary syndrome (HCPS) in animals but target two different antigenic faces on the head domain of Gn. Determining the antigenic sites for neutralizing antibodies will contribute to further therapeutic development for hantavirus-related diseases and inform the design of new broadly protective hantavirus vaccines.

## Main

Orthohantaviruses are emerging zoonotic pathogens that are endemic worldwide and classified into two categories based on the geographic distribution of their reservoir hosts and pathogenesis in humans^1^. Almost sixty virus species have been identified in rodents or shrews, and over twenty species cause disease in humans^2^. Old World hantaviruses (OWHs), including the species Hantaan (HTNV), Puumala (PUUV), Dobrava -Belgrade (DOBV), and Seoul (SEOV), mainly occur in Eastern Europe and China and cause hemorrhagic fever with renal syndrome (HFRS). New World hantaviruses (NWHs), including the species Sin Nombre (SNV) and Andes (ANDV), are endemic in North and South America and cause hantavirus cardiopulmonary syndrome (HCPS). Person-to-person transmission of ANDV has been reported, including in a recent outbreak in Argentina resulting in 34 confirmed cases and 11 fatalities^3, 4^. There are no current FDA-approved medical countermeasures to prevent or treat hantavirus-related disease.

The viral glycoproteins, designated Gn and Gc, form a hetero-tetrameric spike on the surface of the hantavirus virion and facilitate attachment and entry. Previous studies have reported crystal structures for the Gn^5, 6^ and Gc^7, 8^ ectodomains of PUUV and HTNV, and recent work has described the molecular organization of Gn/Gc on the virion surface^9^. Gc is a class II fusion protein and undergoes conformational changes triggered by low pH to mediate the fusion of the viral and host endosomal membranes. Gn is proposed to play a role in receptor attachment and stabilize and prevent the premature fusogenic triggering of Gc^10, 11^. Structural studies have identified a capping loop on Gn that shields the fusion loop on Gc, and the glycoprotein complex is thought to undergo dynamic rearrangements between a closed (or capped) and open (or uncapped) form of the spike^9, 11^.

Recent efforts have identified features of the molecular basis of neutralization by some antibodies targeting HTNV Gn^12^ or PUUV Gc^13^. A rabbit-derived antibody, HTN-Gn1, targets domain A on Gn and overlaps with the putative binding sites for other murine-derived HTNV^14^ and ANDV mAbs^15^. An antibody isolated from a bank vole, P-4G2, targets a site spanning domain I and II on Gc that is occluded in the post-fusion trimeric form, suggesting that the antibody may neutralize through blocking conformational changes required for fusion^8, 13^. Although mAb P-4G2 neutralizes PUUV and ANDV, this activity has only been tested in pseudovirus neutralization assays, and it is unknown if the antigenic site on Gc is occluded on the surface of the authentic virus. Also, these antibodies are not human, and it is unclear what sites are accessible to antibodies during a natural human infection. The first clues towards the sites of vulnerability on the Gn/Gc spike for the human antibody response were recently described by Mittler, Wec, and colleagues^16^. A panel of 135 antibodies were isolated from convalescent PUUV donors, and two distinct neutralizing sites were determined by negative stain electron microscopy (nsEM); one at the Gn/Gc interface and one on prefusion exposed surface of Gc domain I. ADI-42898, a quaternary-site mAb, demonstrated cross-clade neutralizing activity and protected in both PUUV and ANDV hamster post-exposure challenge models. However, mAbs targeting Gn were not described, and, due to its level of surface exposure and variability, the N-terminal domain of Gn likely represents a major site of the neutralizing human antibody response^9, 17^.

Previously, we characterized a panel of human mAbs isolated against Gn/Gc from survivors of SNV or ANDV infection^18^. We demonstrated that NWHs antibodies target at least eight distinct sites on the ANDV Gn/Gc complex, four of which contained potently neutralizing antibody clones, but the location of those sites on the Gn/Gc spike was not known. Here, we define four distinct neutralizing antigenic sites on the Gn/Gc complex. Two potently neutralizing species-specific antibodies, ANDV-5 and ANDV-34, map to two non-overlapping epitopes in the Gn ectodomain and neutralize the virus by blocking attachment. We also show that these antibodies are similar to two human antibody clones previously isolated, MIB22 and JL16^19^. Broadly neutralizing antibodies, ANDV-44 and SNV-53, map to the interface of Gn/Gc, while a less potently neutralizing but broadly reactive antibody, SNV-24, targets domain I of Gc. Both classes of broad antibodies function by inhibiting the triggering of fusion. These data shed light on the basis for both ANDV-specific and broad hantavirus recognition by these antibody clones and suggest that hantavirus vaccine designs should focus on effectively eliciting antibodies to these sites of vulnerability.

## Results

### Neutralizing antibodies target at least four sites on ANDV Gn^H^/Gc spike

The hantavirus glycoprotein spike forms a complex hetero-tetrameric structure on the surface of the virus and is composed of two proteins, Gn and Gc^20^. Gn is separated into two main domains: an N-terminal, membrane-distal Gn head domain (Gn^H^), and a C-terminal Gn base domain (Gn^B^) ^5, 9^. Gn^H^ likely functions in viral attachment, but also has a “capping loop” region that interfaces with the Gc fusion loop to prevent premature exposure of the hydrophobic loop^11^. Due to the surface exposure and sequence variability of Gn^H^, it is likely immunodominant compared to Gc and Gn^B^^17^. Gn^B^ promotes the tetramerization of the complex and is shielded from immune recognition in the Gn/Gc complex.

To determine which subunits of the spike were targeted by neutralizing antibodies, we expressed and purified hantavirus antigens in S2 cells and tested for antibody binding reactivity to the Gn^H^, Gn^B^, or Gc monomeric proteins or the Gn^H^/Gc heterodimer (Fig. 1a). We also generated and purified a “stabilized” form of Gn^H^/Gc by introducing a H953F mutation that prevents the Gc protein from making conformational changes to the post-fusion form^9^. We also produced recombinant IgG1 forms of MIB22 or JL16 (hereafter, rMIB22 or rJL16) based on each antibody’s published heavy or light chain variable genes and cloned them into a human IgG1^19^. All the mAbs tested demonstrated reactivity to the linked ANDV Gn^H^/Gc construct by ELISA, as well as to the “pre-fusion” stabilized form (ANDV Gn^H^/Gc_H953F). Most of the mAbs, apart from ANDV-34 and rMIB22, demonstrated reactivity to Maporal (MAPV) Gn^H^/Gc, a closely related species to ANDV. ANDV nAbs (ANDV-34, ANDV-5, rMIB22, and rJL16) all displayed binding reactivity to Gn^H^, with EC_50_ values less than 100 ng/mL except for rMIB22 (EC_50_ value 5.6 µg/mL). SNV-24 was the only antibody that displayed reactivity to Gc, and we did not detect binding reactivity to Gn^B^ for any mAbs tested. Notably, bnAbs ANDV-44 and SNV-53 did not have detectable binding to Gn^H^, Gn^B^, or Gc alone, and only bound to the linked Gn^H^/Gc antigen, suggesting these antibodies bind a quaternary site only present on the Gn/Gc heterodimer.

**Fig. 1.**
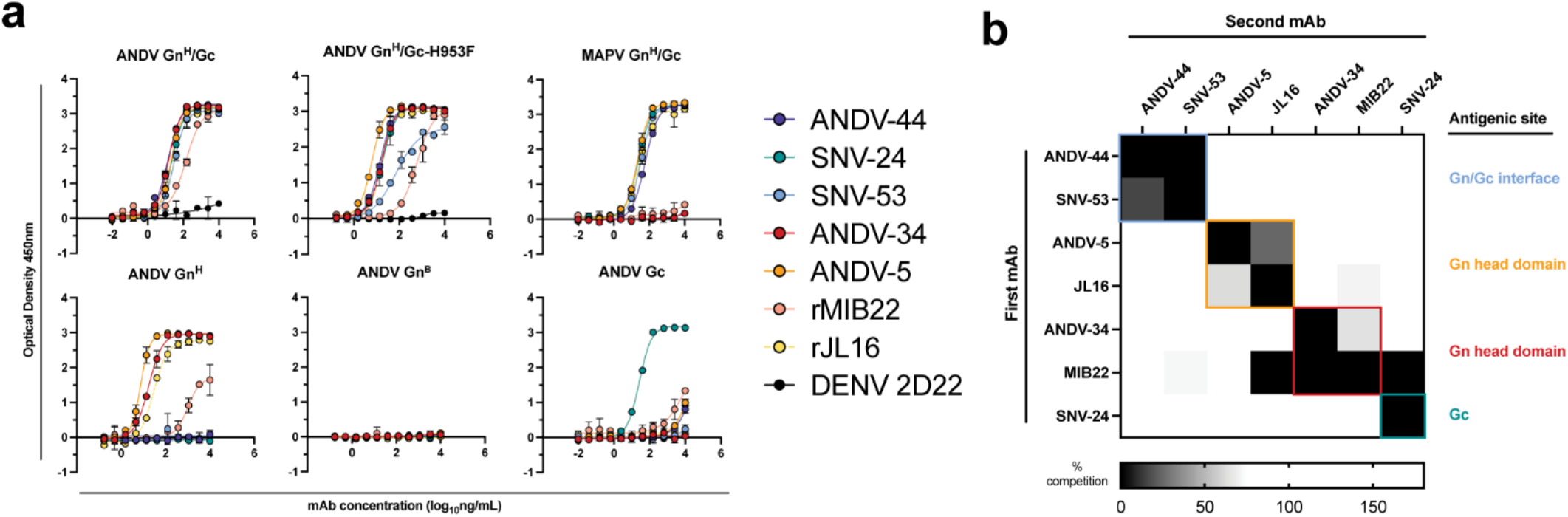
Hantavirus neutralizing antibodies target four distinct regions on the glycoprotein spike. **a.** Binding potency of mAbs to recombinant hantavirus antigens expressed in S2 cells. Binding curves were obtained using non-linear fit analysis, with the bottom of curve constrained to 0, using Prism software. The data shown are representative curves from 3 independent experiments. Mean ± SD of technical duplicates from one experiment are shown. **b.** Competition binding analysis of neutralizing antibodies to ANDV Gn^H^/Gc recombinant protein measured using BLI. % Competition is designated by the heatmap, where black boxes indicate complete competition, gray boxes indicate intermediate competition, and white boxes indicate no competition. The data are shown are representative from 2 independent experiments.

We also performed competition-binding studies utilizing biolayer interferometry and tested the seven mAbs for binding to the linked Gn^H^/Gc antigen (Fig. 1b). As previously shown with a cell surface-displayed version of Gn/Gc, bnAbs ANDV-44 and SNV-53 bin to similar competition groups while SNV-24 bins to a distinct site on Gc. We also determined that rJL16 bins in a group with ANDV-5, while rMIB22 bins with ANDV-34. However, rMIB22 and rJL16 also asymmetrically compete for binding, indicating that all four mAbs likely bind in close spatial proximity to each other on the Gn^H^ domain.

### Low likelihood of viral escape for neutralizing hantavirus antibodies

Determining escape mutations is helpful for designing vaccines and therapeutics to treat hantavirus-related diseases. We next sought to identify neutralization-resistant viral variants for each of the seven nAbs described in this study. To do this, we used a recombinant vesiculostomatitis virus (VSV) pseudotyped with the ANDV or SNV glycoproteins and performed a high throughput escape mutant generation assay using real-time cellular analysis based on similar assays previously described^21–23^. We tested each antibody at saturating neutralization conditions and evaluated escape based on delayed cytopathic effect (CPE) in each individual replicate well. Notably, we did not detect escape mutants using this method for most neutralizing antibodies (*e.g*., SNV-53, ANDV-44, SNV-24, ANDV-34, or ANDV-5) (Fig. 2a). For example, we could not detect any escape mutants through this method for SNV-53, even though we attempted 368 replicates with the VSV/SNV and 184 replicates with the VSV/ANDV viruses for selection (Fig. 2a).

**Fig. 2.**
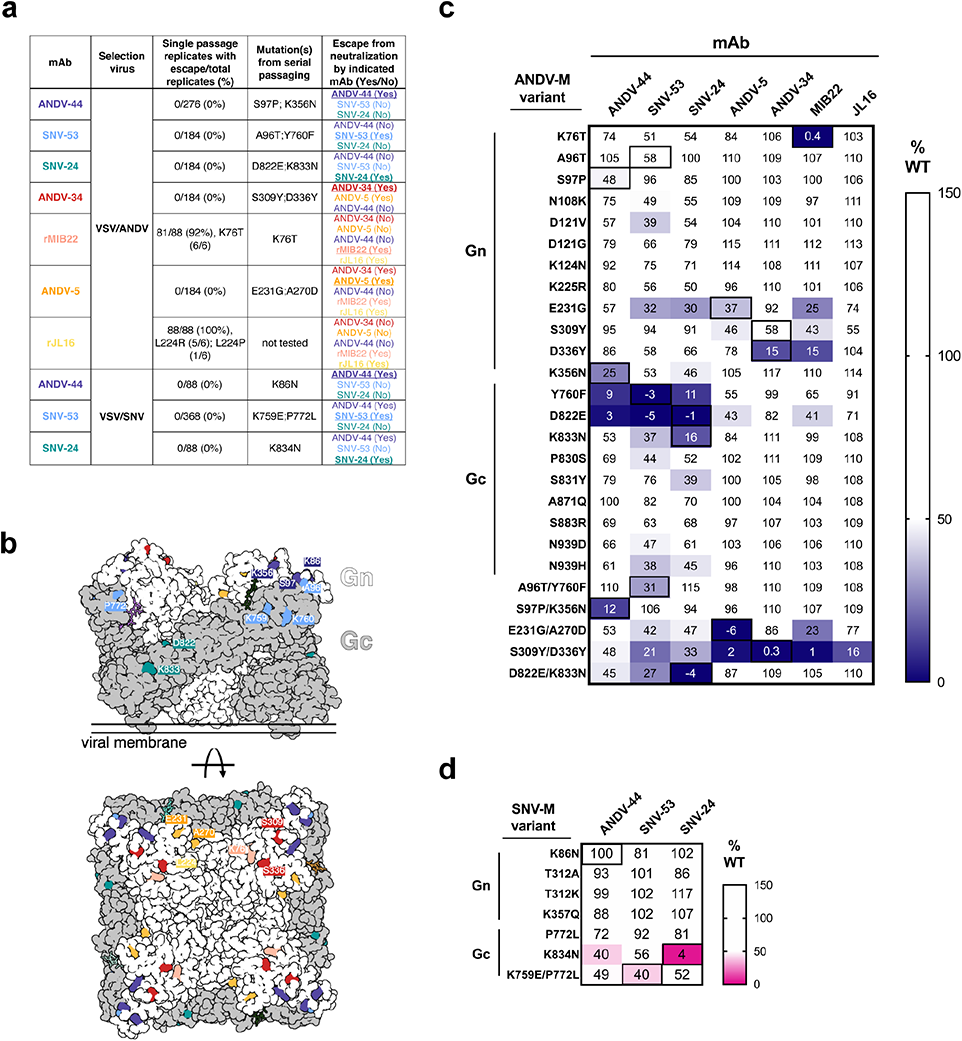
Escape mutant generation and mutagenesis mapping indicate critical binding residues for hantavirus mAbs. **a.** Results from viral escape selection for indicated antibodies. Real-time cellular analysis shows the number of replicates with escape over the total number of replicates for each selection mAb against the indicated selection virus. Mutations from serial passaging were identified for each mAb, and escape was confirmed in the presence of saturating mAb concentrations. VSV/ANDV or VSV/SNV were used for escape selections. **b.** Side and top view of escape mutants mapped to the ANDV Gn/Gc spike (PDB: 6ZJM). The colored spheres designate escape mutants for the indicated antibody. Gn is shown in white, and Gc is shown in grey. **c.** Heatmap of mAb binding in the presence of ANDV mutant constructs. Dark blue boxes indicate loss of binding. The black boxes designate escape mutants for the indicated antibody. The percent binding (% WT) of each mAb to the mutant constructs was compared to the WT SNV or ANDV control. The data are shown as average values from 3-4 independent experiments. All numbering for ANDV sequences was based on GenBank AF291703.2 and SNV sequences were based on GenBank KF537002.1. **d.** Heatmap of SNV mutant constructs as described in Fig. 2c. All numbering for SNV sequences were based on GenBank KF537002.1

In contrast, the CPE profile for 92% and 100% of the replicates for rMIB22 and rJL16, respectively, indicated an escape mutation was present at a high proportion in the viral preparation’s original stock (Fig. 2a). We confirmed resistance to either rMIB22 or rJL16 at 10 µg/mL and sequenced the M-segment of the escaped virus from six replicates. All sequenced rMIB22-selected viruses bore the mutation K76T. Five rJL16-selected virus sequences had an L224R mutation, and one had an L224P mutation. Overall, these results indicate that identifying viral escape to hantavirus antibodies through this method is possible but suggests a low likelihood of *in vitro* escape for potently neutralizing hantavirus nAbs targeting multiple distinct antigenic regions.

### Mapping mutations in antibody escape variant viruses selected with ANDV-specific nAbs or bnAbs

Although we could not select escape mutants in a high-throughput, single passage approach for five of the mAbs, we still wanted to map the critical binding residues and identify escape mutations for the neutralizing antibodies of interest. Thus, we identified escape mutants for mAbs through serial passaging (Fig. 2a).

Potent species-specific neutralizing antibodies selected for mutations located in Gn^H^. ANDV-34-selected viruses contained mutations S309Y and D336Y, two residues on the surface exposed face of Gn domain B (Fig. 2b). rMIB22-resistant variant viruses contained a single mutation, K76T, located in domain A of Gn in close spatial proximity to S336, corresponding with binning analysis for these mAbs (Fig. 2c). Neutralization-resistant viruses selected by ANDV-5 contained mutations E231G and A270D, mapping to α-3 and α-4 helices of the Gn head domain, respectively. Both residues are located near L224, identified in rJL16-resistant viruses, supporting previous data that ANDV-5 and rJL16 compete for a similar binding site (Fig. 1b).

For bnAbs SNV-53 and ANDV-44, we identified mutations in the interface region between Gn/Gc (Fig. 2a). Variant viruses selected by ANDV-44 or SNV-53 contained mutations in both the Gn ectodomain (K356, K86, S97, A96) and Gc domain II near the highly conserved fusion loop (K759, P772, Y760). For SNV-24, we selected escape mutants D822E and K833N in the variant VSV/ANDV viruses and a single homologous mutation, K834N, in the escape-resistant VSV/SNV. Residues K86, Y760, D822, and K833 are highly conserved among members of the *Orthohantavirus* genus (Extended Data Fig. 1).

### Mapping of ANDV-specific nAb and bnAb epitopes by mutagenesis

Since most of the neutralization-resistant viruses contained multiple mutations, we next sought to identify which mutant residues impacted antibody binding. We generated a panel of single- or double-point mutants in the SNV or ANDV M-segment genes based on escape mutants that we selected with mAbs and previously published VSV/ANDV escape mutants^15, 19^. We expressed each mutant M segment gene on the surface of Expi293F cells and used a flow cytometric binding assay to assess how each variant impacted mAb binding. The % wild-type (WT) values were generated by dividing the binding of the mutant by that of the WT construct, and values were normalized based on a positive control oligoclonal mix of mAbs.

The expression levels of SNV and ANDV M-segment variants were comparable to that of the WT constructs, except for three single-point mutants: A270D, C1129F, and K759E (Extended Data Fig. 2). All three mutations were identified in infectious VSVs, so it is not clear why we could not detect the expression of the mutated proteins. Notably, these residues all co-occurred with additional mutations, indicating that these residues may be found in functionally constrained sites.

Most single-point mutants we identified partially impacted the corresponding mAb binding reactivity (Fig. 2c, d). However, the loss-of-binding phenotype was most evident for the double-mutant M-segment genes, suggesting that multiple mutations are required to generate neutralization-resistant variants. Multiple mAbs that mapped to different sites lost binding in the presence of a single mutation. For example, Y760F (a residue near the fusion loop) ablates the binding of ANDV-44/SNV-53 (Gn/Gc interface) and SNV-24 (Gc domain I). Another single point-mutant, D822E (located in Gc domain I), also reduced the binding of all bnAbs. This finding may indicate that this altered residue promotes rearrangements of Gn/Gc that have allosteric effects on interface mAb binding. Additionally, one double mutant, S309Y/D336Y, exhibited a moderate reduction of the binding of the bnAbs and lost binding of ANDV-specific nAbs tested. Both residues are located on Gn domain B and could represent a critical evolutionary strategy to escape species-specific immunity.

### BnAbs recognize two antigenic sites on Gn/Gc

Next, we sought to determine the location of the antigenic sites targeted by bnAbs, SNV-53 and SNV-24, using negative-stain electron microscopy of MAPV Gn^H^/Gc in complex with antigen-binding fragments (Fabs) of antibodies (Fig. 3a). To identify and dock the right Fabs to the maps, we used the information from the binding groups (Fig. 1) and escape mutations (Fig. 2). The 3D reconstructions (EMD-26735) of SNV-53 and SNV-24 in complex with the MAPV Gn^H^/Gc heterodimer showed that the Fabs localize to two distinct regions in proximity to the corresponding viral escape mutations selected (Fig. 3b). SNV-24 bound to Gc domain I, an epitope that is only accessible in the pre-fusion state of Gc and is near the interface between inter-spike Gc heterodimers. P-4G2 (a bank vole-derived mAb)^12^ and group II human PUUV mAbs described previously^16^ also map to a similar site on Gc. This site has limited accessibility in the full virus lattice structure due to neighboring interspike contacts and may results in the incomplete neutralizing activity to authentic virus we demonstrated by these mAbs previously^18^. SNV-53 binds to the capping loop region on Gn, which interfaces with the *cd* and *bc* loops on domain II of Gc (Fig. 3b, 3c). This 3D reconstruction supports our previous finding that bnAbs SNV-53 and ANDV-44 target a quaternary epitope only accessible on the Gn/Gc heterodimer and is consistent with the broadly protective site targeted by ADI-42898 ^16^. Modeling SNV-53 Fab in the context of the (Gn-Gc)_4_ lattice structure shows possible clashes with adjacent spikes in the full virus assembly (Extended Data Fig. 3). However, the low resolution of the model may not fully recapitulate the binding angle and we previously demonstrated that this mAb was capable of complete neutralization of authentic viruses. We also performed hydrogen-deuterium exchange mass spectrometry (HDX-MS) with the Fab forms of SNV-53 or ANDV-44 in complex with ANDV Gn^H^/Gc, and we identified a peptide (residues 81-101) in the capping loop region (Extended Data Fig. 4).

**Fig. 3.**
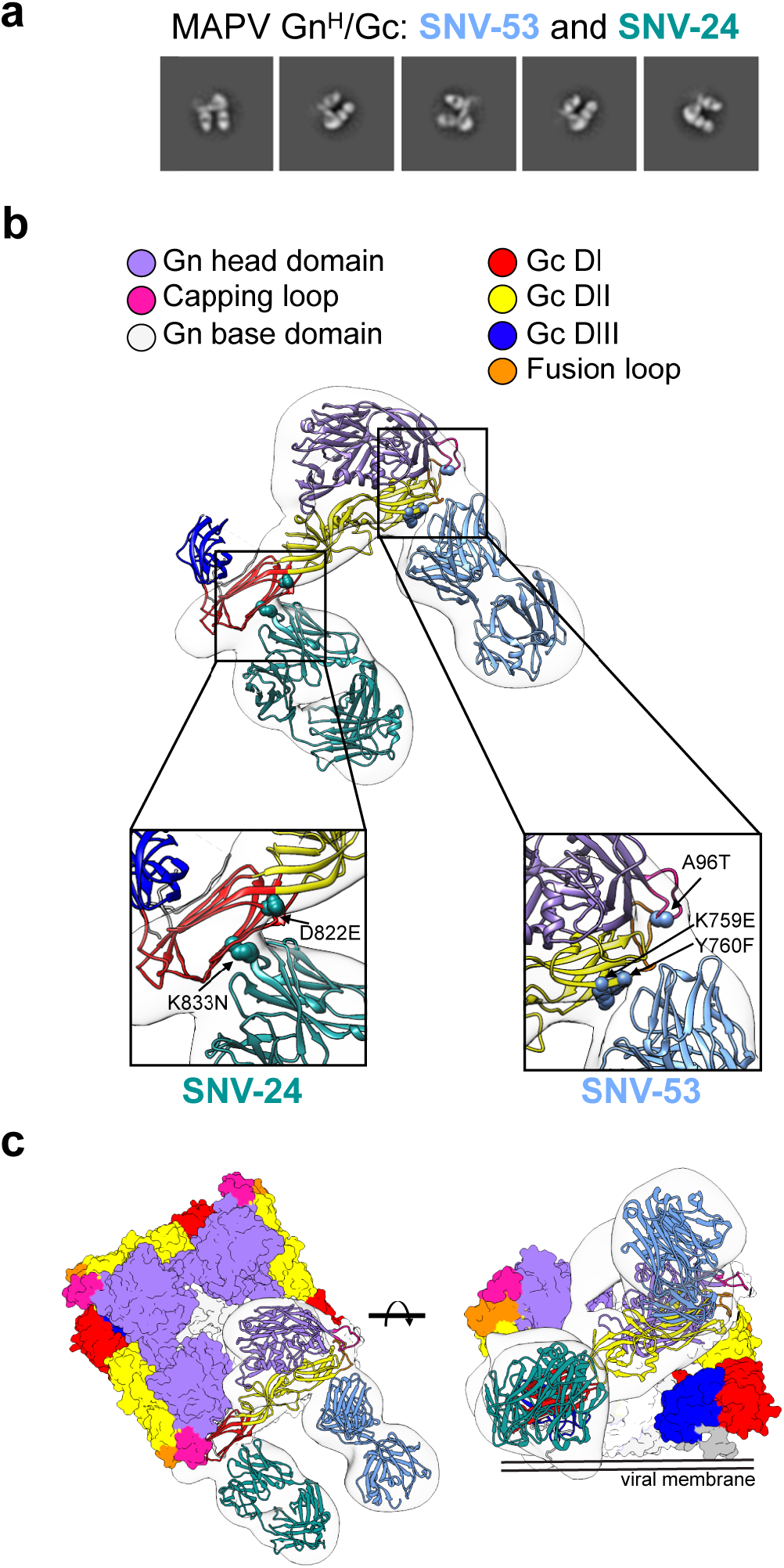
BnAbs SNV-53 and SNV-24 target two sites on the Gn^H^/Gc heterodimer. **a.** Representative nsEM 2D-class averages of SNV-53 and SNV-24 Fabs in complex with MAPV Gn^H^/Gc heterodimer. **b.** Surface representations (light grey) of SNV-53 (blue) and SNV-24 (green) in complex with MAPV Gn^H^/Gc. Escape mutations are indicated by the colored spheres. Gn^H^ is colored in purple, Gn^B^ is colored in light grey, and the capping loop is colored in pink. Domain I, II and III of Gc are colored in red, yellow, and blue, respectively, and the fusion loop is colored in orange. **c.** Model of bnAbs in complex with the (Gn-Gc)_4_ spike as colored in Fig. 3b, top and side view are shown.

Previous sequence analysis determined that SNV-53, which is encoded by human antibody variable region gene segments *IGHV5-51*01/IGLV1-40*01*, is remarkably close to the germline-encoded sequence, with a 97 or 95% identity to the inferred heavy and light chain variable gene sequences, respectively^18^. To understand if somatic mutations are necessary broad recognition of this epitope, we aligned the antibody coding sequence with the inferred germline gene segments and reverted all mutations in the antibody variable regions to the residue encoded by the inferred germline gene. When reverted to its germline form, SNV-53 loses binding activity to all hantavirus species except for SNV (Extended Data Fig. 5a). The germline reverted form of ANDV-44 also has decreased binding activity to ANDV and HTNV, although it has similar reactivity to SNV. SNV-53 also loses all detectable neutralizing activity to both VSV/ANDV and VSV/SNV in its germline reverted form, while ANDV-44 shows decreased potency to VSV/ANDV in its germline-encoded form (Extended Data Fig. 5b). These findings indicate that breadth is developed through affinity maturation, although the critical amino acid changes important for broad reactivity of this quaternary site are unknown.

### Structural basis for neutralization by Gn-targeting antibodies

We were also interested in the epitopes targeted by ANDV-specific mAbs elicited during natural infection, thus, we performed 3D reconstructions of ANDV-5 and ANDV-34 in complex with the ANDV Gn^H^ protomer (EMD-26736), which showed that the mAbs bind to ANDV Gn^H^ at two distinct sites in concordance with the results from competition binning (Fig. 1b). Fab ANDV-5 bound the Gn head domain α3 – α4 angled parallel to the membrane and may interact with a neighbor Gn protomer in the (Gn-Gc)_4_ complex (Fig. 4a). Fab ANDV-34 bound to a distinct site on the opposite face of Gn corresponding to domain B and is angled perpendicular to the viral membrane in the (Gn-Gc)_4_ complex (Fig. 4a).To understand the molecular level interaction of the ANDV-specific neutralizing antibodies, ANDV-5 and ANDV-34, and Gn^H^, we collected a cryo-EM data set and reconstructed a 3D map at 4.1 Å resolution (Fig. 4b and Extended Data Fig. 6, Table 1). The heavy chain solely drives the interactions between ANDV-5 and Gn with four hydrogen bonds and excessive hydrophobic interactions (Fig. 4c, e). The CDRH3 loop interacts with a hydrophobic cavity on Gn (Extended Data Fig. 7). The escape mutation Ala270 is located in the center of the epitope. We could not detect expression of the Gn/Gc glycoproteins bearing the A270D mutation, indicating that A270D may solely ablate ANDV-5 binding but may incur fitness costs offset by mutations (i.e., E231G) that are not located in the epitope. The ANDV-34 interface with Gn is more extensive than the ANDV-5:Gn interface and includes both heavy and light chain interactions with 11 hydrogen bonds (Fig. 4d, f). Both the escape mutations Ser309 and Asp336 are part of the interface, and Ser309 has one hydrogen bond with the epitope.

**Fig. 4.**
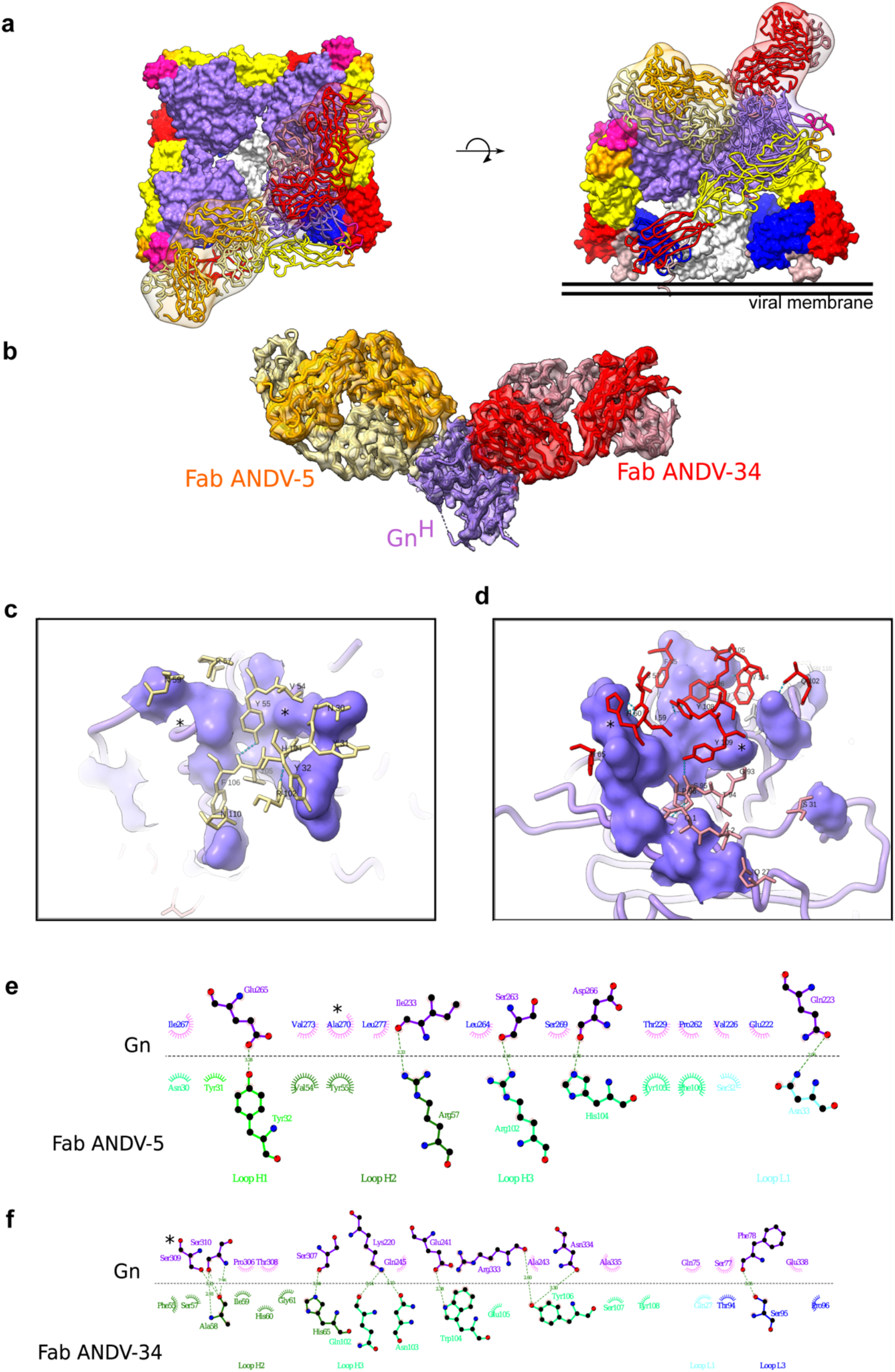
Cryo-EM structure of neutralizing antibodies ANDV-5 and ANDV-34 in complex with ANDV Gn^H^. **a.** Top view (*right*) and side view (*left*) of the heterotetramer Gn/Gc^9^ (PDB: 6ZJM) with low resolution map and model of Gn^(H)^ ANDV-5 and ANDV-34 (purple, orange and red, respectively) complex superimpose with Gn^H^. **b.** Cryo-EM map and model of the Gn-Fabs complex. Right, transparent EM map color by chain with Gn purple, ANDV-5 heavy chain in yellow, ANDV-5 light chain in orange, ANDV-34 heavy chain in red and ANDV-34 light chain in pink. **c.** Zoom-in on the paratope/epitope interface of the ANDV-5:Gn^H^. Fab ANDV-5 residues that is in close contact with Gn. Heavy chain residues (yellow stick), label with single letter and residue number. Gn is shown in purple with the contact residues shown in a surface representation. Asterisks correspond to residues found in the escape mutant viruses. Blue dashed line, H-bond. **d.** Zoom-in on the paratope/epitope interface of the ANDV-34:Gn^H^. Fab ANDV-34 residues that is in close contact with Gn. Heavy residues (red stick), light chain residues (pink stick), label with single letter and residue number. Gn is shown in purple with the contact residues shown in a surface representation. Asterisks correspond to residues found in the escape mutant viruses. Blue dashed line, H-bond. **e.** Fab ANDV-5 paratope and epitope residues involved in hydrogen bonding (dashed lines) and hydrophobic interactions. Hydrophobic interactions residues are shown as curved lines with rays. Atoms shown as circles, with oxygen red, carbon black, and nitrogen blue. Interacting residues that belong to CDR loops are colored in different shade. Image was made with Ligplot+^49^. **f.** Fab ANDV-34 paratope and epitope residues involved in hydrogen bonding (dashed lines) and hydrophobic interactions. Hydrophobic interactions residues are shown as curved lines with rays. Atoms shown as circles, with oxygen red, carbon black, and nitrogen blue. Interacting residues that belong to CDR loops are colored in different shade. Image was made with Ligplot+^49^.

ANDV-5 and ANDV-34 are both encoded by *IGVH1-69* v-genes and use F alleles (*IGHV1-69*01* and \**06*, respectively). Notably, the heavy chain of MIB22 is also encoded by *IGHV1-69*06,* indicating that this gene usage is commonly employed in the neutralizing response found in multiple individuals in response to hantavirus infection. Although all three mAbs are germline-encoded by an F allele at position 55 in the CDHR2 loop, ANDV-34 retains the phenylalanine, but ANDV-5 mutates the residue to a tyrosine, which is a similarly bulky hydrophobic residue, while MIB22 has an F54S mutation (Extended Data Fig. 5e). MIB22 retains the conserved isoleucine at position 54, but ANDV-5 and ANDV-34 mutate the isoleucine to valine, a similarly hydrophobic side chain. Notably, Y55 and V54 make hydrophobic interactions with A270 and neighboring residues. The germline reverted ANDV-5 shows substantially reduced binding to ANDV Gn/Gc and no detectable neutralizing activity to VSV/ANDV (Extended Data Fig. 5c,d). Thus, somatic hypermutation was critical to the potency of ANDV-5, and it is possible that the F55Y/I54V mutation was critical to the interactions that ANDV-5 makes with Gn. MIB22 and ANDV-34 both compete for binding to Gn; thus, *IGHV1-69*06* may encode important germline characteristics that are used to bind to this antigenic site on Gn. F55 hydrophobic interactions with S309, a residue identified in ANDV-34 neutralization resistant viruses. Interestingly, when ANDV-34 is reverted to its germline form, it has a similar binding and neutralization potency as rMIB22 (Extended Data Fig. 5d).

### Neutralization potency is dependent on bivalent interactions

We have previously demonstrated that all antibody clones described here neutralize ANDV or SNV as full-length IgG1 molecules. Here, we sought to determine if bivalency was required for NWH neutralization; therefore, we generated the five antibody clones as recombinant Fab molecules and tested the neutralizing activity. BnAb ANDV-44 retained some neutralizing activity as the Fab form to VSV/ANDV, while SNV-53 Fab form did not have detectable neutralizing activity to VSV/ANDV (Fig. 5a). The opposite was true against VSV/SNV, with SNV-53 demonstrating a decrease in potency, while the ANDV-44 Fab form had no detectable neutralizing activity to VSV/SNV. Gc-specific bnAb SNV-24 only had a slight decrease in neutralization potency in the Fab form. Although it is important to note that we previously demonstrated that SNV-24 has a ∼1,000-fold decrease in IC_50_ neutralization values between the VSV/SNV and the authentic SNV; thus, this finding may not be representative of authentic virus neutralization.

**Fig. 5.**
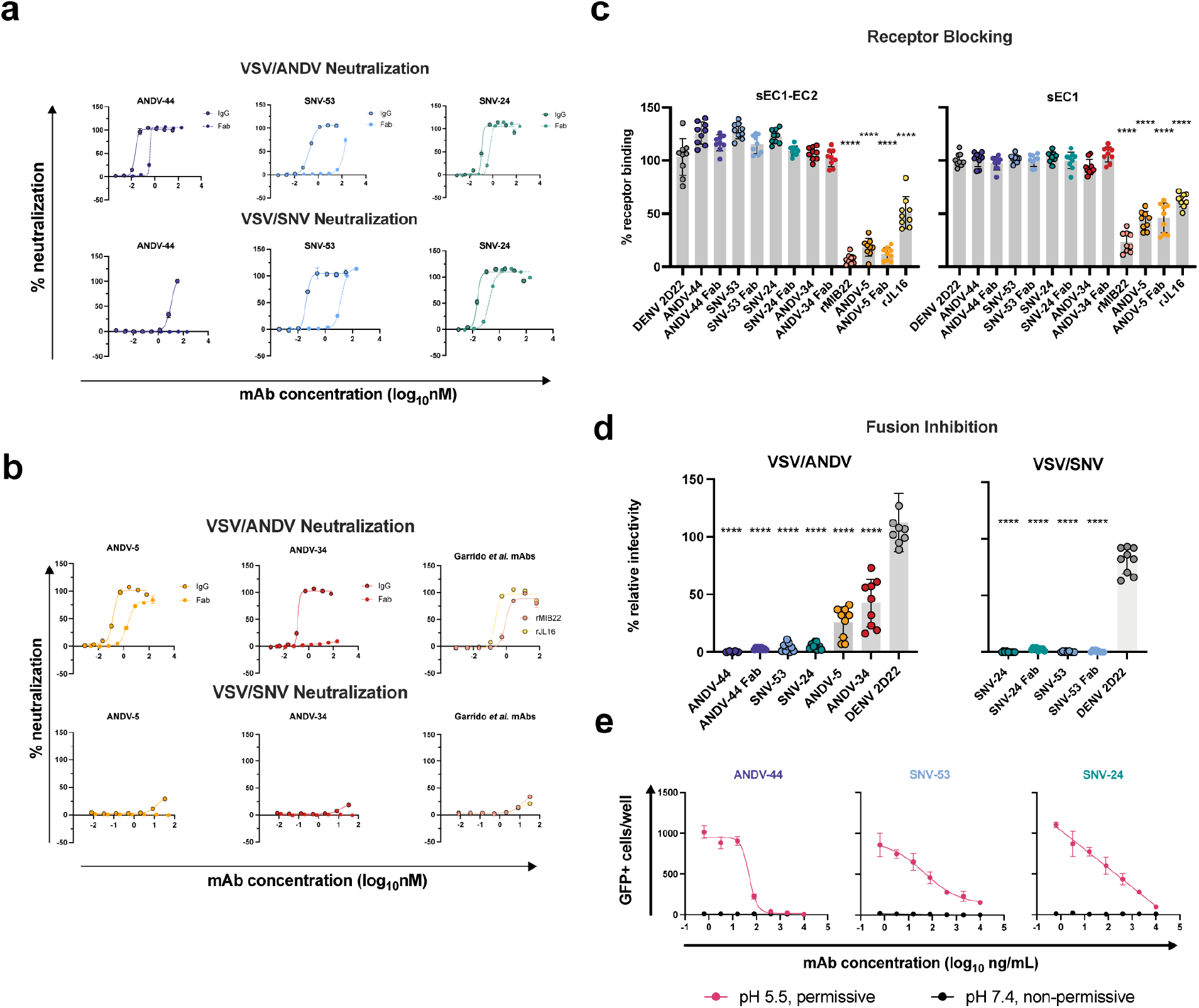
Potently neutralizing hantavirus mAbs inhibit viral entry through viral attachment blocking and/or fusion inhibition. **c.** Neutralization curves of IgG1 and Fab forms of broad mAbs (SNV-53, ANDV-44 or SNV-24) to VSV/SNV and VSV/ANDV determined through real-time cellular analysis. The data shown are representative curves from 3 independent experiments. Mean ± SD of technical duplicates from one experiment are shown. **d.** Neutralization curves of IgG1 and Fab forms of ANDV-nAbs (ANDV-5, ANDV-34, rJL16, or rMIB22) to VSV/SNV and VSV/ANDV determined through real-time cellular analysis. The data shown are representative curves from 3 independent experiments. Mean ± SD of technical duplicates from one experiment are shown. **e.** sEC1 or sEC1-EC2 blocking activity of neutralizing antibodies determined through a flow cytometric assay, in which mAbs were added at saturating concentration before the addition of the each soluble PCDH-1 domain labeled with Alexa Fluor 647 dye. The data shown are averages ± SD from three experiments, n = 9. One-way ANOVA with Dunnett’s test, ****p< 0.0001; ns, non-significant. **f.** FFWO assay testing VSV/ANDV or VSV/SNV post-attachment antibody neutralization in a permissive (pH 5.5) conditions at 30 µg/mL. The data shown are averages ± SD from three experiments, n = 9. One-way ANOVA with Dunnett’s test, ****p< 0.0001. **g.** Representative curves of dose-dependent VSV/ANDV fusion inhibition. The data shown are average values for technical replicates ± SDs. The experiments were performed 3 times independently with similar results.

Gn-specific mAb ANDV-5 retained neutralizing activity as a Fab but showed a slight decrease in neutralization potency (Fig. 5b). We did not detect any neutralizing activity of the Fab form of ANDV-34, despite there being no detectable difference in the binding of Gn^H^ between the IgG1 and Fab form of ANDV-34. This result could indicate that this antibody’s neutralization depends on steric hindrance, not avidity effects. As expected, we did not detect neutralizing activity of ANDV-5, ANDV-34, rMIB22, and rJL-16 to VSV/SNV (Fig. 5b).

### ANDV-specific antibodies neutralize through receptor blocking

New World hantaviruses use PCDH-1 as a receptor for cell attachment^24^. We previously demonstrated that ANDV-5 IgG competes with the extracellular cadherin 1 domain (EC1) of PCDH-1, the primary domain that engages with the Gn/Gc complex. Here, we show that the Fab form of ANDV-5 also blocks both EC1 and EC1-EC2 binding, suggesting that ANDV-5 binds to the receptor-binding site (RBS), although the residues comprising the RBS are currently unknown (Fig. 5c). We also tested mAbs rJL16 and rMIB22 for receptor blocking activity, and both antibodies significantly reduced EC1 and EC1-EC2 binding to ANDV Gn/Gc. Although ANDV-34 does not block EC1 or EC1-EC2 binding to Gn/Gc, it competes for binding to Gn/Gc with rMIB22 and thus likely binds near the receptor-binding site.

### BnAbs neutralize the virus through fusion inhibition

The broadly reactive antibodies did not have detectable receptor-blocking activity in assays with EC1 of PCDH-1, possibly because OWHs do not engage with PCDH-1. Thus, to further explore if the bnAbs neutralize virus at a pre-or post-attachment step in the life cycle, we performed a fusion-from-without (FFWO) assay to test the ability of each antibody to neutralize virus after attachment. All mAbs tested significantly reduced viral infectivity of VSV/ANDV in Vero cell culture monolayers (Fig. 5d,e). However, ANDV-44 completely neutralized VSV/ANDV in the FFWO assay as a full-length IgG1 or Fab molecule. BnAbs SNV-53 and SNV-24 also reduced VSV/ANDV infectivity post-attachment, but both antibodies fully neutralized VSV/SNV as IgG1 or Fab molecules (Fig. 5d). SNV-24 binding maps to the fusogenic protein Gc, while SNV-53 also had critical binding residues located in the capping loop on Gn and the fusion loop on Gc. This result suggests that for the clones studied here, bnAbs target highly conserved fusogenic regions of the Gn/Gc complex and function to neutralize the virus principally through fusion inhibition.

### SNV-53 MAb cross-protects against an Old Word hantavirus when administered prior to exposure

We previously demonstrated that ANDV-44, SNV-53 or SNV-24 provided protection from a lethal ANDV challenge in hamsters^18^. Since SNV-53 potently neutralized HTNV *in vitro*, we next tested if the bnAb could protect hamsters from HTNV infection in an animal prophylaxis model. Syrian golden hamsters are highly susceptible to HTNV infection. Antibodies can be tested in this model for reduction of viral load and prevention of seroconversion after challenge, although the virus does not cause discernable pathological changes in tissues^25–27^. Hamsters were administered 5 mg/kg of SNV-53, SAB-159 (positive control), or DENV 2D22 (negative control) by the i.p. route one day before exposure by the i.m. route with 200 PFU of HTNV (Fig. 6a). The polyclonal human antibody immune globulins SAB-159 were developed through hyperimmunization of transchromosomic bovines with HTNV, and PUUV DNA vaccines, respectively, and previously were demonstrated to protect animals in pre-and post-exposure models of HTNV and PUUV infection^27^. Day 0 bioavailability of neutralizing antibody measured through a pseudovirion neutralization assay (PsVNA) indicated that injected antibody was present in sera as expected for the positive control antibody and SNV-53 (Fig. 6b). PsVNA levels measured on day 28 after virus inoculation revealed that all vehicle-control-treated animals had seroconverted in response to productive infection, while all animals injected with SAB-159 or SNV-53 lacked an increase in PsVNA activity on day 28 (Extended Data Fig. 8a). Day 28 ELISA for antibodies binding to nucleocapsid (N) protein confirmed that animals injected with SAB-159 or SNV-53 did not seroconvert (Fig. 6b).

**Fig. 6.**
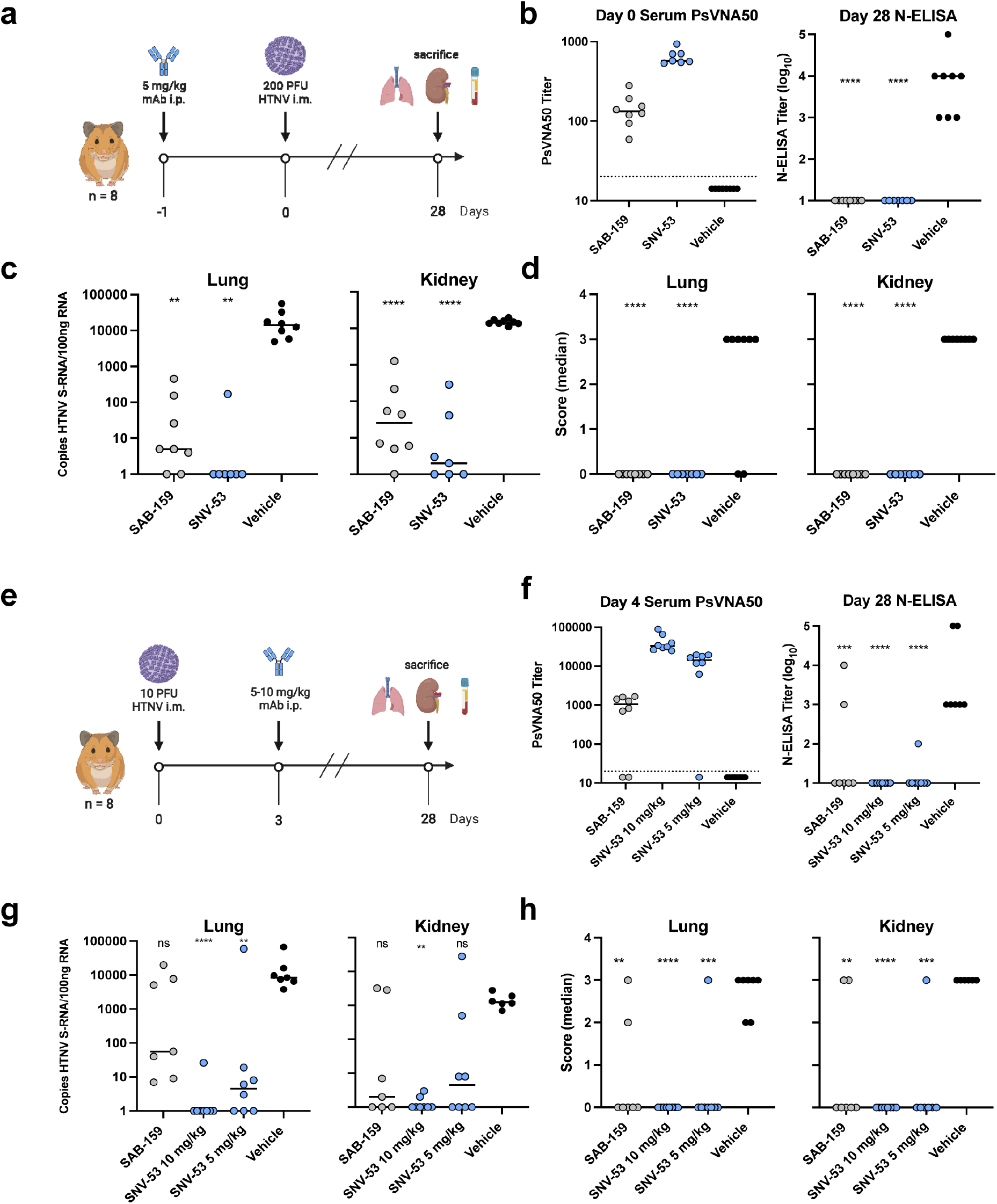
SNV-53 protects hamsters when given before or after HTNV inoculation. **a.** 8-week-old Syrian hamsters (n =8 per treatment group) were administered 5 mg/kg of the indicated Ab treatment and then inoculated 1 day later with 200 PFU of HTNV i.m. All animals were sacrificed at 28 dpi and organs were harvested. **b.** Ab detection in the serum at day 0 by pseudovirion neutralization assay and at day 28 by nucleoprotein ELISA. Dotted line indicated limit of detection. **c.** qRT-PCR detection of HTNV genome in the lungs and kidneys. Kruskal-Wallis test with multiple comparisons of each group to vehicle, * p<0.01, ** p<0.001, *** p<0.001 **** p< 0.0001. ns, not significant. **d.** In-situ hybridization detection of HTNV genome in the lungs and kidneys. One-way ANOVA with multiple comparisons of each group to vehicle, * p<0.01, ** p<0.001, *** p<0.001 **** p< 0.0001. ns, not significant. **e.** 8-week-old Syrian hamsters (n =6 per treatment group) were inoculated with 10 PFU of HTNV i.m., and 5-10 mg/kg of indicated Ab was administered via i.p. route at 3 dpi. All animals were sacrificed at 28 dpi and organs were harvested. **f.** Serum was analyzed as indicated in Fig. 6b. **g.** Lung and kidneys were assayed as described in Fig. 6c. **h.** Lung and kidneys were assayed as described in Fig. 6d.

As expected, pathologic changes in the tissue of the infected hamsters were not detected by hematoxylin and eosin staining; however, a remarkably high amount of virus genome was detected in multiple organs by qRT-PCR and *in situ* hybridization (ISH). Animals injected with SAB-159 or SNV-53 had significantly lower levels of viral RNA in the lungs and kidneys compared to vehicle control-treated animals (Fig. 6c). Lungs and kidneys for all animals injected with SAB-159 or SNV-53 were negative for genome by ISH, whereas all animals injected with vehicle were positive by ISH for genome in both lungs and kidneys (Fig. 6d). Many of the genome-positive organs had large amounts of staining (score 3) despite the presence of neutralizing and anti-N antibodies in the serum that were mostly IgG (Extended Data Fig. 8b,c).

### SNV-53 MAb cross-protects against an Old World hantavirus when administered after virus inoculation

We next tested SNV-53 in a post-exposure model of HTNV infection. Hamsters were inoculated with 10 PFU HTNV by the i.m. route and then treated with SNV-53 at 10 or 5 mg/kg, the positive control SAB-159, or a negative control mAb DENV 2D22 at day 3 post-exposure, and then sacrificed 28 days later and lungs, kidneys, and serum were harvested (Fig. 6e). Day 4 post-infection (1 day after antibody injection) bioavailability studies confirmed presence of SAB-159 or SNV-53 in sera (Fig. 6f). All animals with PsVNA_50_ serum titers lower than ∼200 developed higher serum titers on day 28, indicating the animals had responded to the infection (Extended Data Fig. 8d). Day 28 N-ELISA studies confirmed that post-exposure injection of animals with SAB-159 or SNV-53 significantly reduced seroconversion (Fig. 6f). The most significant impact was when SNV-53 was injected, and there appeared to be a dose-response in which SNV-53 given at a 10 mg/kg dose was most protective. Only 1 of the 16 animals injected with SNV-53 antibody had detectable serum anti-N antibodies on day 28.

qRT-PCR testing detected high levels (>1,000 copies/500 ng RNA) of virus RNA in the lungs of all the control-treated hamsters (Fig. 6g). There was no statistically significant reduction in the RNA levels in any of the treated groups; however, it is notable that the SNV-53 10 mg/kg group had only one of eight animals with a positive signal. There was a similar readout in the kidneys, except that the SNV-53 10 mg/kg protection was statistically significant.

Five of the seven hamsters in the positive control-treated group had no detectable virus RNA as measured by ISH in lungs or kidneys on day 28, which was a significant reduction relative to the vehicle-control-treated group (Fig. 6h). The level of protection for both doses was significant in the lung but not the kidney. All animals injected with the 10 mg/kg dose and all but one animal injected with the 5 mg/mL of SNV-53 had no virus RNA detected in the lungs or kidneys. The one positive animal in the SNV-53 with a high level of virus in both lung and kidney was the same animal that did not receive the correct antibody dose as measured PsVNA on day-4 sera (Fig. 6f).

### ANDV-specific antibodies protect from ANDV challenge in hamsters

We previously demonstrated that ANDV-5 was partially protective against lethal disease in hamsters after ANDV infection ^18^. Previous studies have shown that inoculation of Syrian golden hamsters with ANDV elicits a pathological response like that seen during HCPS in humans^28^. To achieve complete protection and test ANDV-34 for therapeutic efficacy in this model, Syrian golden hamsters were inoculated with 200 PFU of ANDV virus (strain Chile-9717869) and administered 10 mg/kg of ANDV-5, ANDV-34, or DENV-2D22 (isotype control) on days 2 and 5 post-inoculation (Fig. 7a). Both ANDV-5 and ANDV-34 demonstrated complete protection from disease and mortality, whereas all but one of the isotype control-treated animals succumbed on day 9 (Fig. 7b). Thus, antibodies elicited to both antigenic sites on Gn^H^ can provide protection from lethal HCPS-like disease *in vivo*.

**Fig. 7.**
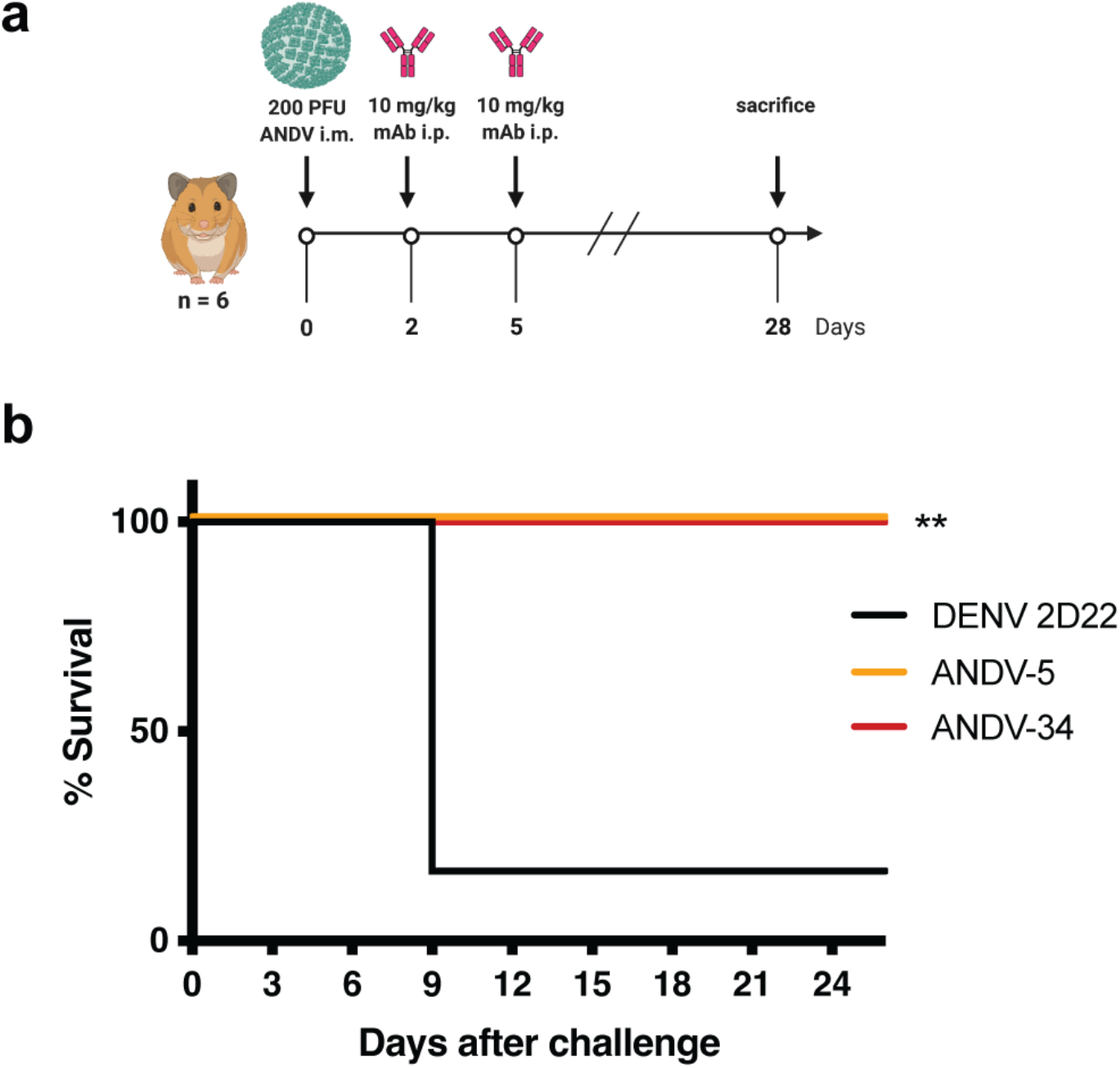
ANDV-34 and ANDV-5 protect Syrian golden hamsters for lethal ANDV challenge. **a.** 8-week-old Syrian hamsters (n =6 per treatment group) were inoculated with 200 PFU of ANDV i.m., and 10 mg/kg of indicated mAb was administered via i.p. route at 2 and 5 dpi. Animals were treated with a dengue-specific mAb (DENV 2D22) to serve as an isotype control. **b.** Kaplan-Meier survival plot. Statistical analysis of survival curves was done using a log-rank (Mantel-Cox) test comparing each group to the control (DENV 2D22), **, p=0.0014.

## Discussion

Antibodies isolated from survivors of hantavirus infection have been shown to reduce viral replication and protect against disease in animal models^16, 18, 19^, suggesting that mAbs might be effective for preventing or treating hantavirus-related disease in humans. The selection of optimal lead candidate antibodies for monotherapy or rational composition of an antibody combination requires detailed information on the antigenic sites of vulnerability to inhibition. Here, we defined four major antigenic sites on the hantavirus Gn/Gc complex recognized by neutralizing and protective antibodies. The antibodies are promising for development as biologics, and the landscape analysis of protective sites that emerges from the mapping studies provides a blueprint for virus species-specific or broadly protective vaccine design.

Two classes of broad antibodies we describe exhibit breadth and likely neutralize through viral fusion inhibition. These antibodies are strikingly similar to group I and group II bnAbs isolated from PUUV convalescent donors^16^, demonstrating the immunodominance of these antigenic sites recognized by the human humoral repertoire. Apart from flaviviruses, canonical class II fusions proteins are hidden in the three-dimensional structure of the glycoprotein spike by an accompanying protein to prevent premature fusion during viral entry^29^. Once viral attachment occurs, the virion is taken up in the endosome, where low-pH triggers the dissociation of the complex and exposes the hydrophobic fusion loop initiating viral-host cell fusion and entry into the cytoplasm. We propose that one class of hantavirus bnAbs we have studied here (i.e., SNV-53 and ANDV-44) stabilize the association of Gn (the accompanying protein) to Gc (the fusion protein) by binding to and cross-linking the two subunits – preventing the heterodimer from undergoing conformational changes necessary for fusion.

BnAbs that span the heterodimeric interface also have been described for other enveloped viruses, including hepatitis C virus (HCV)^30^, Rift Valley Fever virus (RVFV)^31^, and chikungunya virus^32^. One antibody, RVFV-140, isolated from a Rift Valley Fever virus survivor, potently neutralizes RVFV through a fusion inhibition mechanism and targets an epitope only present on the Gn/Gc hetero-hexamer^31^. Interface targeting antibodies can aid in the structural determination of heterodimers; the structure of the hepatitis C virus E1E2 heterodimer was determined by co-expressing the complex with a bnAb, AR4A, which targets an epitope proximal to the E1E2 interface^33^. Altogether, the data indicate that there exists an essential class of bnAbs that target quaternary epitopes at the interface regions on heterodimers for many viruses. These antibodies appear to be induced by viruses in diverse families that use heterodimeric surface glycoproteins, and the shared mechanism appears to be fusion inhibition by preventing the complex conformational changes required for viral fusion. The structures of the antigenic sites recognized by these rare antibodies could provide a blueprint for structure-based reverse vaccinology efforts to design stabilized viral immunogens to elicit a broad, protective antibody response^34^.

SNV-53 did cross-protect both in prophylactic and therapeutic small animal models of HTNV and ANDV infection, suggesting that SNV-53 is a promising candidate for treating both HCPS and HFRS. Based on studies using germline revertant forms of these antibodies, SNV-53 and ANDV-44 have developed their breadth and neutralization potency across multiple species through the accumulation of somatic mutations. It may be possible to continue to affinity mature SNV-53-like antibodies to increase the potency and breadth across species. Combined with the recent description of Gn/Gc interface mAbs from PUUV convalescent donors^16^, our findings indicate that these types of antibodies can be readily elicited during natural infection and are likely critical to pan-hantavirus therapeutic and vaccine development.

The second class of bnAbs, exemplified by SNV-24, targets the domain I region on Gc. This region is highly conserved, which likely explains why SNV-24 demonstrates broad reactivity. The putative epitope for SNV-24 is near the epitope identified for bank vole-derived antibodies 1C9 and 4G2, and a putative mechanism for this antibody class is the inhibition of viral fusion through blocking the formation of a post-fusion trimer^13, 35^. However, this site contacts neighboring Gc protomers, therefore, it may not be fully accessible on the lattice virion surface. These accessibility considerations may contribute to the observed incomplete neutralization of authentic hantaviruses by SNV-24^18^. SNV-24 does potently neutralize hantavirus glycoprotein pseudotyped VSVs, but this finding might be due to an alternative arrangement of the glycoprotein spikes on the VSV virion that does not correctly mimic the glycoprotein array on authentic viruses.

Although understanding bnAbs is important, ANDV-specific nAbs will still be critical to deploying medical countermeasures. ANDV represents the only hantavirus shown to transmit person-to-person and causes notable outbreaks yearly in Argentina and Chile^4^. Potently neutralizing ANDV antibodies – represented by ANDV-5 and ANDV-34 – target two distinct faces of the globular, membrane distal Gn head domain. The epitope for ANDV-5 maps to the α-3 and α-4 helices, while ANDV-34 targets domain B. Both sites of vulnerability do not overlap with the epitopes determined for previously described Gn-targeting mAbs, HTN-Gn1 and nnHTN-Gn2^12^. A canonical neutralization mechanism used by potent antibodies is to block the attachment of viruses to host cells. Here, we demonstrate that selected ANDV-neutralizing human mAbs block virus binding to the receptor for hantavirus entry protocadherin-1 (PCDH-1), specifically to the first extracellular (EC-1) domain amino-acid (residues 61–172) or EC-2 domain (residues 173–284) ^24^. The finding that ANDV-specific, potently neutralizing mAbs function through PCDH-1 blockade is consistent with the finding that PCDH-1 is used by NWHs but not OWHs. Although the RBS on hantavirus glycoproteins is currently unknown, we propose that the site likely overlaps with the epitope of ANDV-5 since the Fab form of ANDV-5 also competes with sEC1-EC2 indicating that steric clashes are the not the primary driver of receptor blocking. However, it also possible that the RBS spans two Gn:Gn protomers, as we demonstrated that ANDV-5 Fab may contact neighboring, intraspike Gn subunits. Both antibodies protect against HCPS-like disease in Syrian hamsters, indicating that both epitopes on the Gn can elicit a protective neutralizing antibody response.

We also show that JL16 can be categorized in a similar antibody class as ANDV-5, while MIB22 can be classified into a similar class as ANDV-34. Notably, three ANDV-specific nAbs (ANDV-5, ANDV-34, and MIB22) from two different human donors are encoded by *IGHV1-69* and use the F alleles *\*01* and *\*06*. *IGHV1-69* encoded antibodies are prevalent in response to many different pathogens, including influenza^36^, HIV^37^, HCV^38^, EBOV^39^ and even *Staph aureus*^40^. V_H_1-69 gene usage is overrepresented in antiviral antibodies due to the genetic characteristics of the germline-encoded CDRH2 loop^41^. Rational vaccine design should promote the formation of V_H_1-69 antibodies since they have a low somatic hypermutation barrier to high-affinity binding and can be readily elicited in most individuals. Hydrophobic interactions facilitated by the templated amino acids in the CDHR2 loop may contribute to recognizing ANDV Gn by naïve B cells during acute infection.

RNA viruses are particularly prone to mutations and rapid viral evolution, therefore, prospective identification of escape mutations and defining the likelihood of escape is an essential step in developing antibody combinations. Strikingly, five of the nAbs tested here were particularly resistant to escape, and most escape mutants eventually selected through serial passaging had multiple mutations. This finding could indicate that a single mutation is insufficient to confer antibody escape while maintaining the functionality required for viral fitness. Although it was challenging to generate neutralization-resistant variants, we did show that some single mutations could completely ablate the binding of multiple antibodies, even if the mutation was located distant from the putative binding site in the three-dimensional structure. It has been previously demonstrated in other viruses, such as SARS-CoV-2, that single mutations can have functional consequences on the 3D structure of antigenic proteins, thereby altering antibody activity. For example, S371L confers resistance to numerous SARS-CoV-2 antibodies from multiple classes^42^, while E406W causes allosteric remodeling of the receptor-binding domain that significantly impacts the binding of the REGEN-CoV therapeutic monoclonal antibody cocktail^43, 44^. Using antibody combinations that resist escape from single-point mutations is an important objective. MIB22 and JL16 have demonstrated efficacy as a monotherapy and a combination therapy *in vivo*^19, 45^. It is not clear the benefit of combining these two mAbs, since they partially compete for binding, have a similar mechanism of action, and are vulnerable to escape from similar amino acid changes (S309Y/D336Y).

There is still much to learn about the humoral immune response to the hantavirus Gn and Gc proteins, including where major functional epitopes are. Here, we begin to unravel the epitopes targeted by neutralizing antibodies elicited during infection with New World hantaviruses. This endeavor is critical for vaccine design against numerous pathogens, including RSV^46^, HIV^47^, and, more recently, SARS-CoV-2^48^. To prepare for the possible emergence of novel hantaviruses, it is imperative to understand the functional antigenic sites on the glycoprotein spike that need to be presented to the immune response to generate an effective vaccine.

## Acknowledgments

We thank Dr. Rachael Wolters, Dr. Robert Carnahan, Dr. Seth Zost, Dr. Naveen Suryadevara, and Dr. Pavlo Gilchuk for intellectual contributions and general experimental support, and Ryan Irving, Alex Bunnell, Rachel Nargi and Brandon Somerville for technical, managerial and project management support. The recombinant VSV/SNV and VSV/ANDV reagents were a kind gift from K. Chandran and R. Jangra. We thank the UTMB Animal Resource Center for the support of the animal study. We thank Mary L. Milazzo for facilitating BSL-3 and ABSL-4 training and transferring viral isolates. The work of T.B.E. was supported by NIH training grant 5T32GM008320-30. The work of J.W.H. and other USAMRIID authors was supported by the Military Infectious Disease Research Program under project number MI210048. Opinions, interpretations, conclusions, and recommendations are those of the author and are not necessarily endorsed by the U.S. Army. EM data collections were conducted at the Center for Structural Biology Cryo-EM Facility at Vanderbilt University. We acknowledge the use of the Glacios cryo-TEM, which was acquired by NIH grant 1 S10 OD030292-01.

## Author Contributions

Conceptualization, T.B.E., J.W.H., and J.E.C.; Methodology, T.B.E., R.L.B., N.A.K., A.B. J.W.H., and J.E.C.; Investigation, T.B.E., R.L.B., N.A.K., E.B., L.M.P., B.S., S.A.K., R.K.K., N.S.C., M.S.P., L.S.H., S.M., I.A.Z., J.X.R., A.T.; Writing – Original Draft, T.B.E., E.B.. R.L.B., J.W.H., and J.E.C.; Writing –Review & Editing, T.B.E., R.L.B., N.A.K., A.B., J.W.H. and J.E.C.; Funding Acquisition J.W.H., A.B. and J.E.C.; Supervision, A.B., J.W.H., and J.E.C.

## RESOURCE AVAILABILITY

### LEAD CONTACT

Further information and requests for resources and reagents should be directed to and will be fulfilled by the Lead Contact, James E. Crowe, Jr.

### MATERIALS AVAILABILITY

Materials described in this paper are available for distribution for nonprofit use using templated documents from Association of University Technology Managers “Toolkit MTAs”, available at: https://autm.net/surveys-and-tools/agreements/material-transfer-agreements/mta-toolkit.

### DATA AND CODE AVAILABILITY

All data needed to evaluate the conclusions in the paper are present in the paper or the Supplemental Information. The reagents used in this study are available by Material Transfer Agreement with Vanderbilt University Medical Center. Original/source data for all Figs. in the paper is available at Mendeley Data.

## MATERIALS AND METHODS

### Cell lines

Expi293F cells (Thermo Fisher Scientific, Cat# A14527; RRID:CVCL D615, female) were cultured in suspension at 37°C in 8% CO_2_ shaking at 125 RPM in Freestyle F17 Expression Medium (GIBCO Cat# A13835-01) supplemented with 10% Pluronic F-68 and 200 mM of L-glutamine. ExpiCHO cells (Thermo Fisher Scientific Cat# A29127; RRID:CVCL 5J31, female) were cultured in suspension at 37°C in 8.0% CO_2_ shaking at 125 RPM in ExpiCHO Expression Medium (Thermo Fisher Scientific). Vero cell lines (ATCC,Cat# CCL-81, RRID:CVCL_0059, African green monkey, female) were cultured at 37°C in 5% CO_2_ in DMEM (Thermo Fisher) supplemented with 10% fetal bovine serum. *Drosophila* S2 cells (Thermo Fisher Scientific Cat# 51-4003) were cultured in suspension at 37°C shaking at 125 RPM in Schneider’s *Drosophila* Medium (Thermo Fisher Scientific, Cat# 21720001) supplemented with 10% fetal bovine serum. All cell lines were tested for mycoplasma monthly, and all samples were negative.

### Viruses

Replication-competent, recombinant VSV strains bearing ANDV or SNV glycoproteins were kindly provided by K. Chandran and propagated on Vero cells as previously described ^24^. HTNV 76-118 ^50^ was plaque purified twice and passaged in Vero E6 cells, as previously described ^26^. Andes virus strain Chile-9717869 (Chile R123) were obtained from the World Reference Center for Emerging Viruses and Arboviruses housed at UTMB.

### Recombinant human IgG1 and Fab expression and purification

ExpiCHO cells were transiently transfected using the ExpiCHO Expression System (GIBCO) with plasmids encoding human IgG1 or Fab cDNAs. Supernatant was harvested from ExpiCHO cultures and filtered with 0.45-mm pore size filter flasks. HiTrap MabSelectSure columns (Cytiva) or CaptureSelectTM CH1-XL columns (Thermo Fisher Scientific) were used to affinity purify IgG1 or Fab from ExpiCHO supernatant using an ÄKTA pure protein purification system (Cytiva).

### Expression and purification of hantavirus antigens

Soluble, recombinant hantavirus antigens were expressed and purified as previously described^9^. Genes coding for the ANDV Gn head domain (GenBank: NP_604472.1, residues 21-374) and the Gc ectodomain (GenBank: NP_604472.1, residues 652–1107) were combined into a single chain using a flexible linker (GGSGLVPRGSGGGSGGGSWSHPQFEKGGGTGGGTLVPRGSGTGG), and single domain ANDV constructs for Gn base domain (Gn^B^, residues 375-484), Gn head (Gn^H^, residues 21– 374), and Gc (residues 652–1107) were also generated. A similar linked construct was made for Maporal virus (MAPV, strain HV-97021050, NCBI code YP_009362281.1). The constructs were codon-optimized for *Drosophila* cell expression and cloned into a plasmid (pT350) containing an MT promoter, BiP signal sequence, and C-terminal double strep tag (GenScript, kindly provided by P. Guardado-Calvo and F. Rey). Drosophila S2 cells were transfected with the pT350-ANDV Gn^H^/Gc and pCoBlast (Thermo Fisher Scientific) plasmids at a 19:1 ratio, respectively, and stably-transfected cells were selected using 25 µg/mL of blasticidin. Stable cell lines were maintained in Schneider’s *Drosophila* Medium supplemented with 25 µg/mL blasticidin and grown in shaker flasks to a density of 1 x 10^7^ cells/mL and induced with 4 µM CdCl_2_. S2 cell supernatant was collected after 5 days and supplemented with 10 µg/mL of avidin and purified through a StrepTrap HP column (Cytiva). The sample was purified further by size-exclusion chromatography on HiLoad 16/600 Superdex column (Cytiva).

### Negative-stain electron microscopy

Electron microscopy imaging was performed, as previously described^51, 52^, with ANDV Gn^H^ protein in complex with ANDV-5 and ANDV-34 and MAPV Gn^H^/Gc in complex with SNV-53 and SNV-24. Recombinant forms of ANDV Gn^H^ and MAPV Gn^H/^Gc were expressed and purified as described above. Fabs forms of ANDV-5, ANDV-34, SNV-53, and SNV-24 were expressed and purified as described above. Complexes were generated by incubating the recombinant proteins with the two corresponding Fabs in a 1:1.2:1.2 (antigen:Fab:Fab) molar ratio. 3 µL of the sample at ∼10 µg/mL was applied to a glow-discharged grid with continuous carbon film on 400 square mesh copper electron microscopy grids (Electron Microscopy Sciences). Grids were stained with 2% uranylformate^53^. Images were recorded on a Gatan US4000 4kX4k CCD camera using an FEI TF20 (TFS) transmission electron microscope operated at 200 keV and control with SerialEM^54^. All images were taken at 50,000 magnification with a pixel size of 2.18 Å/pixel in low-dose mode at a defocus of 1.5 to 1.8 mm. The total dose for the micrographs was 33 e/Å^2^. Image processing was performed using the cryoSPARC software package^55^. Images were imported, CTF-estimated and particles were picked automatically. The particles were extracted with a box size of 180 pix and binned to 96 pix (4.0875 Å/pixel) and multiple rounds of 2D class averages were performed to achieve clean datasets. The final dataset was used to generate Initial 3D volume and the volume was refine for final map at the resolution of ∼18 Å. Model docking to the EM map was done in Chimera^56^. For the ANDV Gn^H/^Gc protomer or tetramer PDB: 6Y5F or 6ZJM were used and for the Fab PDB:12E8. Chimera or ChimeraX software was used to make all Figures, and the EM map has been deposited into EMDB (EMD-26735 and EMD-26736).

### Cryo-EM sample preparation and data collection

Two Fabs ANDV-5, ANDV-34 and ANDV Gn^H^ were expressed recombinantly and combined in a molar ration of 1:1.2:1.2 (Ag:Fab:Fab). The mixture was incubated for over-night at 4°C and purified by gel filtration. 2.2 µl of the purified mixture at concentration of 0.2 mg/mL was applied to glow discharged (40 s at 25mA) UltraAu grid (300 mesh 1.2/1.3, Quantifoil). The grids were blotted for 3 s before plunging into liquid ethane using Vitrobot MK4 (TFS) at 20°C and 100% RH. Grids were screened and imaged on a Glacios (TFS) microscope operated at 200 keV equipped with a Falcon 4 (TFS) DED detector using counting mode and EEF. Movies were collected at nominal magnification of 130,000×, pixel size of 0.73 Å/pixel and defocus range of 0.8 to 1.8 µm. Grids were exposed at ∼1.18 e/aÅ^2^/frame resulting in total dose of ∼50 e/Å^2^ (Extended Fig. 5 **and** Table 1)

### Cryo-EM data processing

Data processing was performed with Relion 4.0 beta2^57^. EER Movies were preprocessed with Relion Motioncor2^58^ and CTFFind4^59^. Micrographs with low resolution, high astigmatism and defocus were removed from the data set. The data set was autopicked first by Relion LoG ^60^ and was subject to 2D classification. Good classes were selected and used for another round of autopicking with Topaz training and Topaz picking ^61^. The particles were extracted in a box size of 180 pixel and binned to 128 pixels (pixel size of 2.053 Å/pixel). The particles were subjected to multiple rounds of 2D class averages, 3D initial map and 3D classification to obtain a clean homogeneous particle set. This set was re-extracted at a pixel size of 1.46 Å/pixel and was subjected to 3D autorefinement. The data were further processed with CTFrefine ^62^, polished and subjected to final 3D autorefinement and postprocessing. Detailed statistics are provided in Extended Fig. 5 **and** Table 1.

### Model building and refinement

For model building PDB: 6Y5F^9^ was used for Gn^H^. Three AlphaFlod2 predictions for the Fv of ANDV-5, Fv of ANDV-34 or Fc segment of the Fab was used for the Fabs. All the models were first docked to the map with Chimera^56^ or ChimeraX^63^. The Gn^H^ and the Fc were rigid-body real-space refined with Phenix^64^. To improve the coordinates of ANDV-5 and ANDV-34 Fv, the models were subjected to iterative refinement of manual building in Coot^65^ and Phenix. The models were validated with Molprobity^66^ (Extended Data Table 1).

### Bio-layer interferometry competition binding analysis

An Octet Red96 instrument (Sartorius) was used for competition binding analysis of antibodies. Octet® Streptavidin (SA) Biosensors (Sartorius, Cat # 18-5019) were incubated in 200 µL of 1× kinetics buffer (diluted in D-PBS) for 10 minutes. Baseline measurements were taken in 1× kinetics buffer for 60 seconds and then recombinant ANDV Gn^H^/Gc (10 µg/mL) was loaded onto the tips for 180 seconds. The tips were washed for 30 seconds in 1× kinetics buffer and then the first antibody (50 µg/mL) was associated for 300 seconds. The tips were washed again for 30 seconds in 1× kinetics buffer and then the second antibody (50 µg/mL) was associated for 300 seconds. The maximal binding of each antibody was calculated by normalizing to the buffer only control. The Octet epitope software was used to determine the percent competition for each antibody and antibodies were clustered using the Pearson correlation.

### ELISA binding assays

384-well plates were coated with 1 µg/mL of purified recombinant protein overnight at 4°C in carbonate buffer (0.03 M NaHCO_3_, 0.01 M Na_2_CO_3_, pH 9.7). Plates were incubated with blocking buffer (2% non-fat dry milk, 2% goat serum in PBS-T) for 1 hour at room temperature. Serially-diluted primary antibodies were added to appropriate wells and incubated for 1 hour at room temperature, and horseradish peroxidase-conjugated Goat Anti-Human IgG (Southern Biotech, Cat# 2040-05, RRID:AB_2795644) were used to detect binding. TMB substrate (Thermo Fisher Scientific) was added, and plates were developed for 5 minutes before quenching with 1N hydrochloric acid and reading absorbance at 450 nm on a BioTek microplate reader.

### Escape mutant generation

VSV/ANDV or VSV/SNV escape mutants were generated by serially passaging the viruses in Vero cell monolayer cultures with increasing concentrations of SNV-53, SNV-24, or ANDV-44. VSV/ANDV or VSV/SNV (MOI of ∼1) were incubated for 1 hour at 37°C with each antibody at their respective IC_50_ value, and the mixture was added to Vero cells in a 12-well plate. Cells were monitored for CPE and GFP expression 72 hours after inoculation to confirm virus propagation, and 200 µL of supernatant containing the selected virus was passaged to a fresh Vero monolayer in the presence of increasing concentrations of the selection antibody. After 6 to 7 passages, escape mutant viruses were propagated in a 6-well culture plate in the presence of 10 µg/mL of the selection antibody. Viruses were also propagated in the presence of 10 µg/mL of the non-selection antibodies to assess susceptibility. Viral RNA was then isolated using a QiAmp Viral RNA extraction kit (QIAGEN) from the supernatant containing the selected viral mutant population. The ANDV or SNV M gene cDNA was amplified with a SuperScript IV One-Step RT-PCR kit (Thermo Fisher Scientific) using primers flanking the M gene (Forward primer: 5′-ATGCTGCAATCGTGCTACAA-3′; Reverse primer: 5′-GACTGCCGCCTCTGCTC-3′). The resulting amplicon (∼3,500 bp) was purified using SPRI magnetic beads (Beckman Coulter) at a 1:1 volume ratio and sequenced by the Sanger sequence technique using primers giving forward and reverse reads of the entire M segment (VSV/ANDV primers: 5’-GCTGTGTTACGATCTGACTTGC-3’, 5’-CGGGGTGCCCATGTATTCAT-3’, 5’-AGCCTTGCCGACTGATTTGA-3’, 5’-CCAGGCTCCCTCTATTGCTT-3’; VSV/SNV primers: 5’-AGCCTGTAATCAAACACATTGTCT-3’, 5’-GGCATTGCCTTTGCTGGAG-3’, 5’-ACACACCGCAAGCAGTACAT-3’, 5’-CCCATGCTCCGTCGATACTT-3’). The full-length M gene segments for the original VSV/SNV and VSV/ANDV stock used was also confirmed through Sanger sequencing. Each read was aligned to the VSV/ANDV or VSV/SNV genome to identify mutations.

### Site-directed mutagenesis of M segment gene

Plasmids containing a cDNA encoding the full-length M segment from SNV (pWRG/SN-M(opt) ^67^ and pWRG/AND-M(opt2) ^68^ were used mutagenized using the Q5 Site-Directed Mutagenesis Kit (NEB, Cat# E0554S). Primers were designed using the NEBaseChanger tool (https://nebasechanger.neb.com/), and PCR reactions were performed according to the manufacturer’s manual. Plasmids were transformed into NEB^®^ 5-alpha Competent *E. coli* cells (New England Biolabs; Cat# C2987I). All mutants were sequenced using Sanger DNA sequencing to confirm the indicated nucleotide changes.

### Expression of mutant M segments

For expression of mutant constructs, we performed micro-scale transfection (700 µL) of Expi293F cell cultures using the Gibco ExpiFectamine 293 Transfection Kit (Thermo Fischer Scientific, Cat# A14525) and a protocol for deep 96-well blocks (Nest, Cat# 503162). Briefly, sequence confirmed plasmid DNA was diluted in OptiMEM I serum-free medium and incubated for 20 minutes with ExpiFectamine 293 Reagent (Gibco). The DNA– ExpiFectamine 293 Reagent complexes were used to transfect Expi293F cell cultures in 96-deep-well blocks. Cells were incubated, shaking at 1,000– 1,500 rpm inside a humidified 37°C tissue culture incubator in 8% CO_2_ and harvested 48 hours post-transfection.

### Flow cytometric binding analysis to mutant M segments

A flow cytometric assay was used to screen for and quantify binding of antibodies to the mutant constructs. Transfected Expi293F were plated into 96-well V-bottom plates at 50,000 cells/well in FACS buffer (2% ultra-low IgG FBS, 1 mM EDTA, D-PBS). Hantavirus-specific mAbs or an irrelevant mAb negative control, rDENV-2D22, were diluted to 1 µg/mL in FACS buffer and incubated 1 hour at 4°C, washed twice with FACS buffer, and stained with a 1:1,000 solution of goat anti-human IgG antibodies conjugated to phycoerythrin (PE) (Southern Biotech, Cat# 2040-09; RRID:AB_2795648) at 4°C for 30 minutes. Cells then were washed twice with FACS buffer and stained for 5 minutes at 4°C with 0.5 mg/mL of 40,6-diamidino-2-phenylindole (DAPI) (Thermo Fisher). PE and DAPI staining were measured with an iQue Screener Plus flow cytometer (Intellicyt) and quantified using the manufacturer’s ForeCyt software. A value for percent PE-positive cells was determined by gating based on the relative fluorescence intensity of mock transfected cells. Tab

### Real-time cell analysis (RTCA) neutralization assay

A high-throughput RTCA assay that quantifies virus-induced cytopathic effect (CPE) was used to test antibody-mediate virus neutralization under BSL-2 conditions, using a general approach previously described for other viruses ^21^. VSV/ANDV and VSV/SNV were titrated by RTCA on Vero cell culture monolayers to determine the concentration that elicited complete cytopathic effect (CPE) at 36 hours post-inoculation. 50 µL of Dulbecco’s Modified Eagle Medium (DMEM) supplemented with 2% FBS was added to each well of a 96-well E-plate (Agilent) to establish the background reading, then 50 µL of Vero cells in suspension (18,000 cells/well) were seeded into each well to adhere. Plates were incubated at RT for 20 minutes, and then placed on the xCELLigence RTCA analyzer (formerly ACEA Biosciences, now Agilent). Cellular impedance was measured every 15 minutes. VSV/ANDV (∼1,000 infectious units [IU] per well) or VSV/SNV (∼1,200 IU per well) were mixed 1:1 with mAb in duplicate in a total volume of 100 µL using DMEM supplemented with 2% FBS and incubated for 1 hour at 37°C in 5% CO_2_. At 16 to 18 hours after seeding the cells, the virus-mAb mixtures were added to the cells. Wells containing virus only (no mAb added) or cells only (no virus or mAb added) were included as controls. Plates were measured every 15 min for 48 hours post-inoculation to assess for inhibition of CPE as a marker of virus neutralization. Cellular index (CI) values at the endpoint (36 hours after virus inoculation) were determined using the RTCA software version 2.1.0 (Agilent). Percent neutralization was calculated as the CI in the presence of mAb divided by the cells only (no-CPE control) wells. Background was subtracted using the virus only (maximum CPE) control wells. Half maximal inhibitory concentration (IC_50_) values were calculated by nonlinear regression analysis using Prism software version 9 (GraphPad).

### RTCA escape mutant screening

The RTCA method described above was modified to generate escape mutant viruses after a single passage under saturating neutralizing antibody concentrations ^22, 23, 69^. 0.5, 1, 5, 10, or 20 µg/mL (depending on the potency of the antibody to the given virus) of the selection antibody was mixed 1:1 in 2% FBS-supplemented DMEM with VSV/ANDV (∼10,000 IU per well) or VSV/SNV (∼12,000 IU per well) and incubated for 1 hour at 37°C and then added to cells. Virus only (no mAb) and medium only (no virus) wells were included as controls. Escape mutant viruses were identified by a drop in cellular impedance over 96 hours. Supernatants from wells containing neutralization resistant viruses were expanded to 12-well plates in saturating concentrations of the selection antibody to confirm escape-resistant phenotype and the presence of other neutralizing mAbs as a control. Supernatants were filtered and stored at −80°C.

### Hydrogen-deuterium exchange mass spectrometry (HDX-MS)

ANDV GnH-Gc protein and Fabs were prepared at 15 pmol/µL. Labeling was done in Dulbecco’s phosphate-buffered saline (DPBS), pH 7.4, in H_2_O, pH 7.4 (no labelling) or D_2_O, pH 7.0 (labelling). Samples were incubated for 0, 10, 100, 1,000 or 5,000 seconds at 20°C. The labelling reaction was quenched by addition of 50 µL TCEP quench buffer, pH 2.4, at 0°C. Automated HDX incubations, quenches, and injections were performed using an HDX-specialized nano-ACQUITY UPLC ultraperformance liquid chromatography (UPLC) system coupled to a Xevo G2-XS mass spectrometer. Online digestion was performed at 20°C and 11.600 psi at a flow of 150 µL/min of 0.1% formic acid in H_2_O (3.700 psi at 40 µL/min), using an immobilized-pepsin column. Peptides were identified in un-deuterated samples using Waters ProteinLynx Global Server 3.0.3 software with non-specific proteases, minimum fragment ion matches per peptide of three and oxidation of methionine as a variable modification. Deuterium uptake was calculated and compared to the non-deuterated sample using DynamX 3.0 software. Criteria were set to a minimum intensity of 5,000, minimum products 4 and a sequence length 5 to 25 residues.

### Fusion from without (FFWO) assay

A FFWO assay that measures fusion-dependent antibody neutralization was used as previously described for other bunyaviruses ^31^. Sterile, cell-culture-treated 96-well plates were coated overnight at 4°C with 50 µg/mL of poly-d-lysine (Thermo Fisher Scientific). Coated plates were washed with D-PBS and dried, and Vero cells were plated at 18,000 cells/well and incubated at 37°C in 5.0% CO_2_ for 16 hours. The supernatant was aspirated, and cells were washed with binding buffer (RPMI 1640, 0.2% BSA, 10 mM HEPES pH 7.4, and 20 mM NH_4_Cl) and incubated at 4°C for 15 minutes. VSV/ANDV was diluted to an MOI of ∼1 in binding buffer and added to cells for 45 minutes at 4°C. Cells were washed to remove unbound viral particles. Hantavirus-specific or control mAbs were diluted in DMEM supplemented with 2% FBS and added to cells for 30 minutes at 4°C. Fusion buffer (RPMI 1640, 0.2% BSA, 10 mM HEPES, and 30 mM succinic acid at pH 5.5) was added to cells for 2 minutes at 37 °C to induce viral FFWO. To measure pH-independent plasma membrane fusion of viral particles, control wells were incubated with RPMI 1640 supplemented with 0.2% BSA and 10 mM HEPES (pH 7.4) for 2 minutes at 37°C. The medium was aspirated, and cells were incubated at 37°C in DMEM supplemented with 5% FBS, 10 mM HEPES, and 20 mM NH_4_Cl (pH 7.4). After 16 hours, the medium was removed, the cells were imaged on an CTL ImmunoSpot S6 Analyzer (CTL), and GFP-positive cells were counted in each well.

### HTNV hamster challenge and passive transfer

Female Syrian hamsters (*Mesocricetus auratus*) 6 to 8 weeks of age (Envigo, Indianapolis, IN) were anesthetized by inhalation of vaporized isoflurane using an IMPAC6 veterinary anesthesia machine. Once anesthetized, hamsters were injected with the indicated concentration of HTNV diluted in 0.2 mL PBS by the i.m. (caudal thigh) route using a 25-gauge, 1-inch needle at a single injection site. Antibodies were administered at the indicated concentrations and days to anesthetized hamsters by i.p. injection using a 23-gauge, 1-inch needle. Vena cava blood draws occurred on anesthetized hamsters within approved blood collection limitations. Terminal blood collection occurred under KAX (ketamine-acepromazine-xylazine) anesthesia and prior to pentobarbital sodium for euthanasia. All animal procedures were conducted in an animal biosafety level (ABSL-3) laboratory.

### HTNV pseudovirion neurtralization assay (PsVNA)

The PsVNA using a non-replicating VSVΔG-luciferase pseudovirion system was performed as previously described^70^. The plasmid used to produce the HTNV pseudovirions was pWRG/HTN-M(co)^71^.

### Nucleoprotein (N) ELISA

The ELISA used to detect N-specific antibodies (N-ELISA) was described previously^72, 73^.

### Isolation of RNA and real-time PCR

RNA isolation from homogenized lung and kidney tissues and real-time PCR were conducted as previously described^74^.

### *In situ* hybridization tissue studies

Lung and kidney tissue sections were fixed, stained, and mounted as previously described^75^. Slides were scored by intensity ranging from 0 to 3, with median values presented (Extended Data Fig. 7c).

### ANDV hamster challenge

Animal challenge studies were conducted in the ABSL-4 facility of the Galveston National Laboratory. The animal protocol for testing of mAbs in hamsters was approved by the Institutional Animal Care and Use Committee (IACUC) of the University of Texas Medical Branch at Galveston (UTMB) (protocol #1912091). 8-week-old female golden Syrian hamsters (*Mesocricetus auratus*) (Envigo) were inoculated with 200 PFU of Andes hantavirus (strain Chile-9717869) by intramuscular (i.m.) route on day 0. Animals (n = 6 per group) were treated with 10 mg/kg of rANDV-34, rANDV-5, or rDENV 2D22 (a negative control dengue virus-specific antibody) by the i.p. route on days 2 and 5 after virus inoculation. Body weight and body temperature were measured each day, starting at day 0. On day 28 post-challenge, all animals were euthanized with an overdose of anesthetic (isoflurane or ketamine/xylazine) followed by bilateral thoracotomy.

## QUANTIFICATION AND STATISTICAL ANALYSIS

Statistical analysis for each experiment is described in Methods and/or in the Figure legends. All statistical analysis was done in Prism v7 (GraphPad) or RStudio v1.3.1073.

### Binding and neutralization curves

EC_50_ values for mAb binding were calculated through a log transformation of antibody concentration using four-parameter dose-response nonlinear regression analysis with a bottom constraint value of zero. IC_50_ values for mAb-mediated neutralization were calculated through a log transformation of antibody concentration using four-parameter dose-response nonlinear regression analysis with a bottom constraint value of zero and a top constraint value of 100. EC_50_ and IC_50,_ values were generated from 2 to 3 independent experiments and reported as an average value.

### Testing of protective efficacy in Syrian hamsters

Studies were done with 6-8 hamsters per antibody treatment group. Survival curves were created through Kaplan-Meier analysis, and a log-rank (Mantel-Cox) test was used to compare each mAb treatment group to the isotype control (rDENV 2D22) (* p < 0.05, ** p < 0.01, *** p < 0.001, **** p<0.0001)

### N-ELISA and PsVNA

Data analyzed using a one-way analysis of variance (ANOVA) with multiple comparisons for experiments with groups of ≤ 3 groups. In all analyses, a *P* value of <0.05 was considered statistically significant.

## Extended Data Figures and Figure Legends

**Extended Data Fig. 1.**
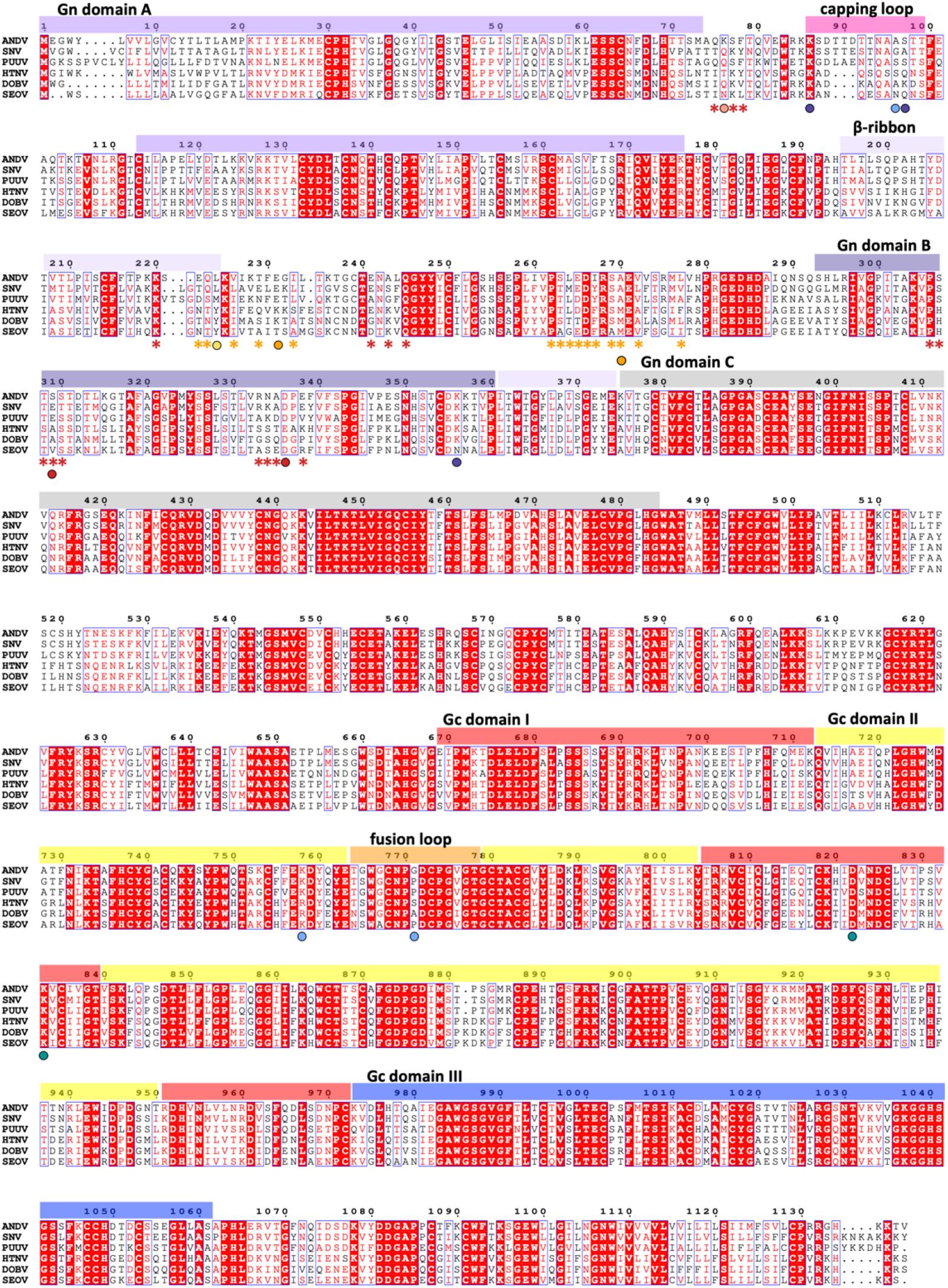
Amino acid alignment of hantavirus species. Multiple sequence alignment of the M-segment from six representative Orthohantaviruses. Domains are colored as followed, Gn domain A: purple, domain B: dark purple, β-ribbon domain: light purple, insertions in white, capping loop: pink; Gn domain C in grey. Gc: domain I: red, domain II: yellow, domain III: dark blue, fusion loop: orange. Strictly conserved residues are highlighted in a red background, and similar residues are colored red. Escape mutations are indicated by colored spheres; SNV-53: light blue, SNV-24: green, ANDV-44: indigo, ANDV-5: orange, rJL16: yellow, ANDV-34: red, rMIB22: salmon. Contact residues for ANDV-5 and ANDV-34 are indicated by orange or red asterisks, respectively. Sequences were aligned from Andes virus (ANDV, NC_003467.2), Sin Nombre virus (SNV, L37903.1), (Puumala virus, (PUUV, KJ994777.1), Hantaan virus (HTNV, JQ083394.1), Dobrava-Belgrade virus (DOBV, JF920149.1), and Seoul virus (SEOV, NC_005237.1). Figure was generated using ESprit 3.0 ^1^.

**Extended Data Fig. 2.**
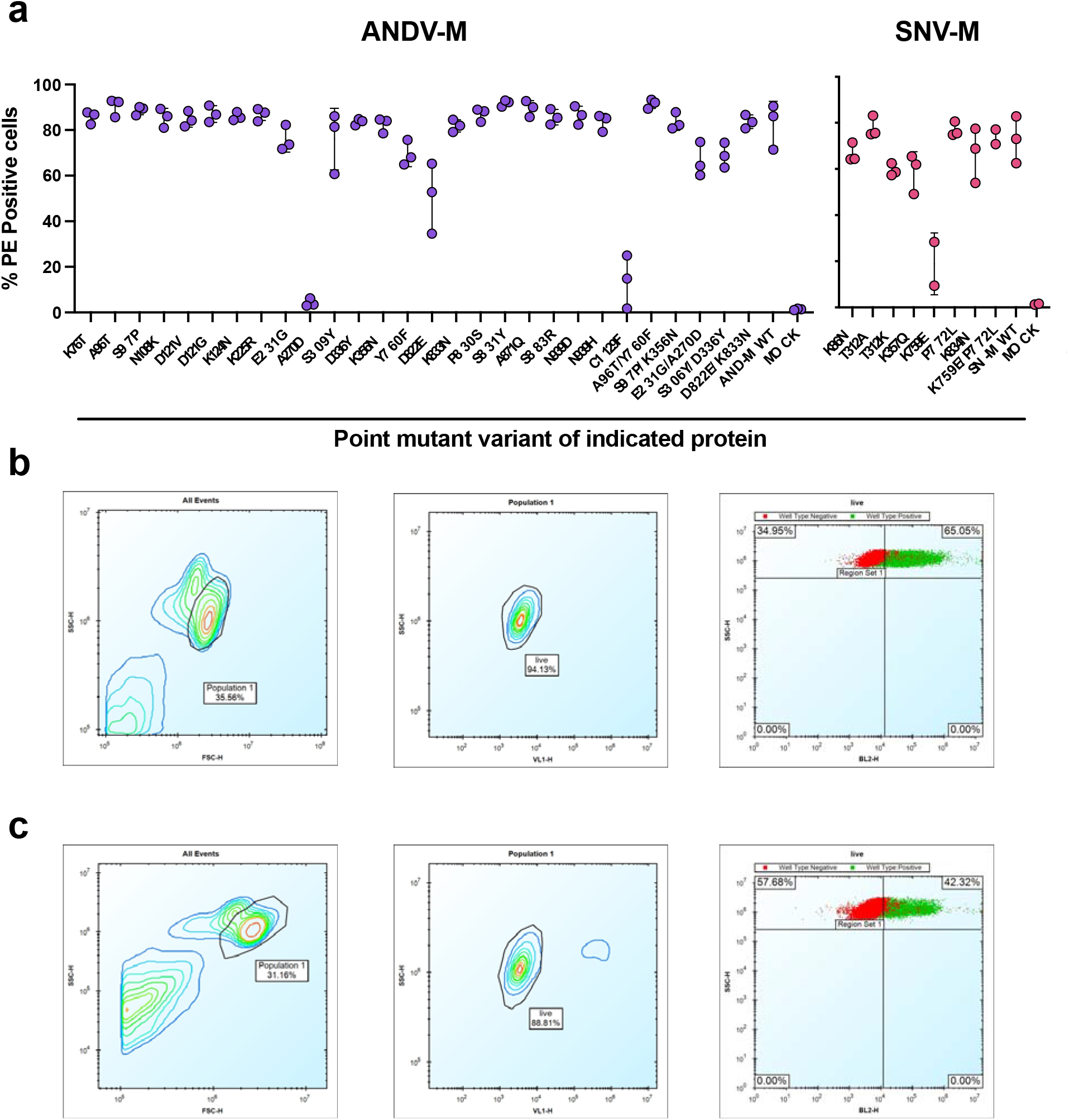
Mutagenesis expression levels and gating strategy. **a.** Expression levels of Gn/Gc point mutants were measured based on binding of a positive control oligoclonal mix of antibodies targeting multiple epitopes on the Gn/Gc antigens. The value for % PE+ cells was determined by gating on untransfected Expi293F cells (mock). The data are shown as average values from 2-3 independent experiments. **b.** Gating strategy for antibody binding to Expi293F cells expressing ANDV M-segment. Cells were first gated by forward and side scatter and dead cells were excluded using a viability dye (DAPI). Binding of the positive control sample (oligoclonal mix of hantavirus-reactive antibodies), shown in green, was detected with a PE-conjugated goat anti-human IgG secondary. The gate for the glycoprotein-specific subset (PE+) was placed based on staining of untransfected Expi293F cells, shown in red. **c.** Gating strategy for antibody binding to Expi293F cells expressing SNV M-segment, as described in Extended data fig. 2b.

**Extended Data Fig. 3.**
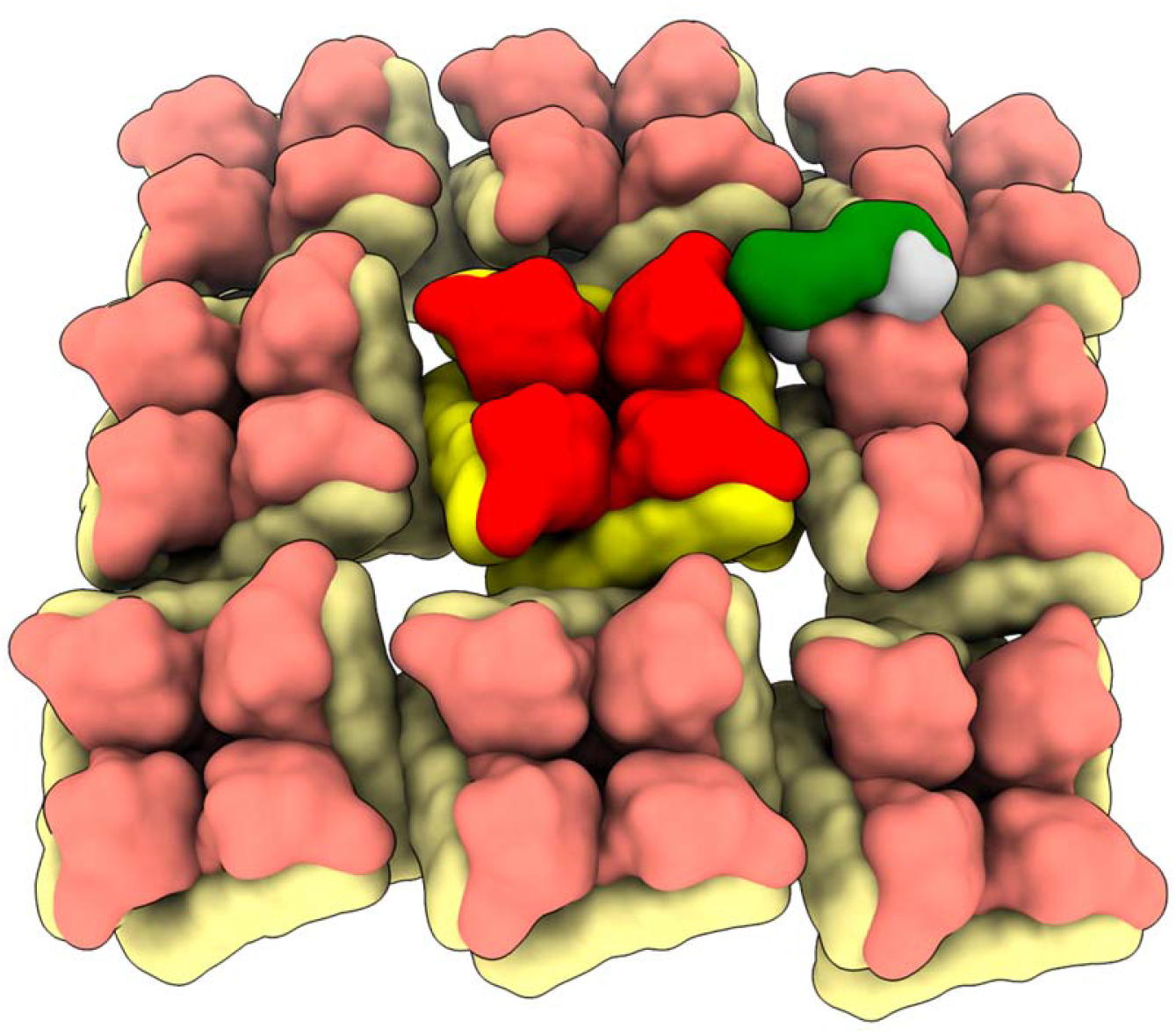
SNV-53 Fab docked to the hantavirus surface glycoprotein lattice (EMD-11236). Model is colored as follows, Gn: red, Gc: yellow, SNV-53 Fab: green/blue.

**Extended Data Fig. 4.**
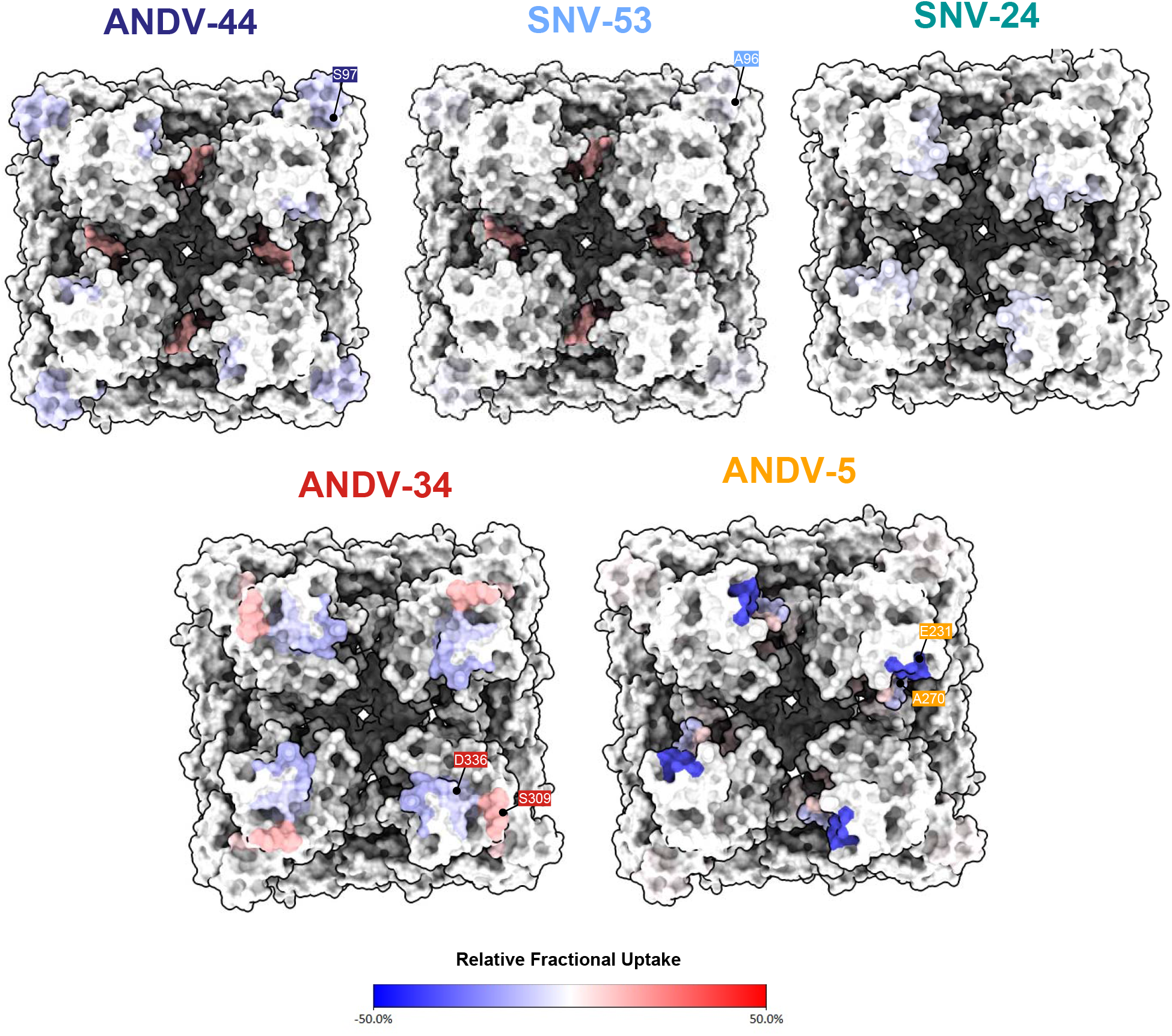
Hydrogen-deuterium exchange mass spectrometry analysis of hantavirus antibodies in complex with ANDV Gn^H^/Gc. Relative fractional deuterium uptake difference at the 10,000 s time point was mapped onto the cryo-EM reconstruction of the glycoprotein complex (PDB: 6ZJM). The relative fractional uptake difference (%) is colored from blue to red. Escape mutants located in the identified peptides are labeled.

**Extended Data Fig. 5.**
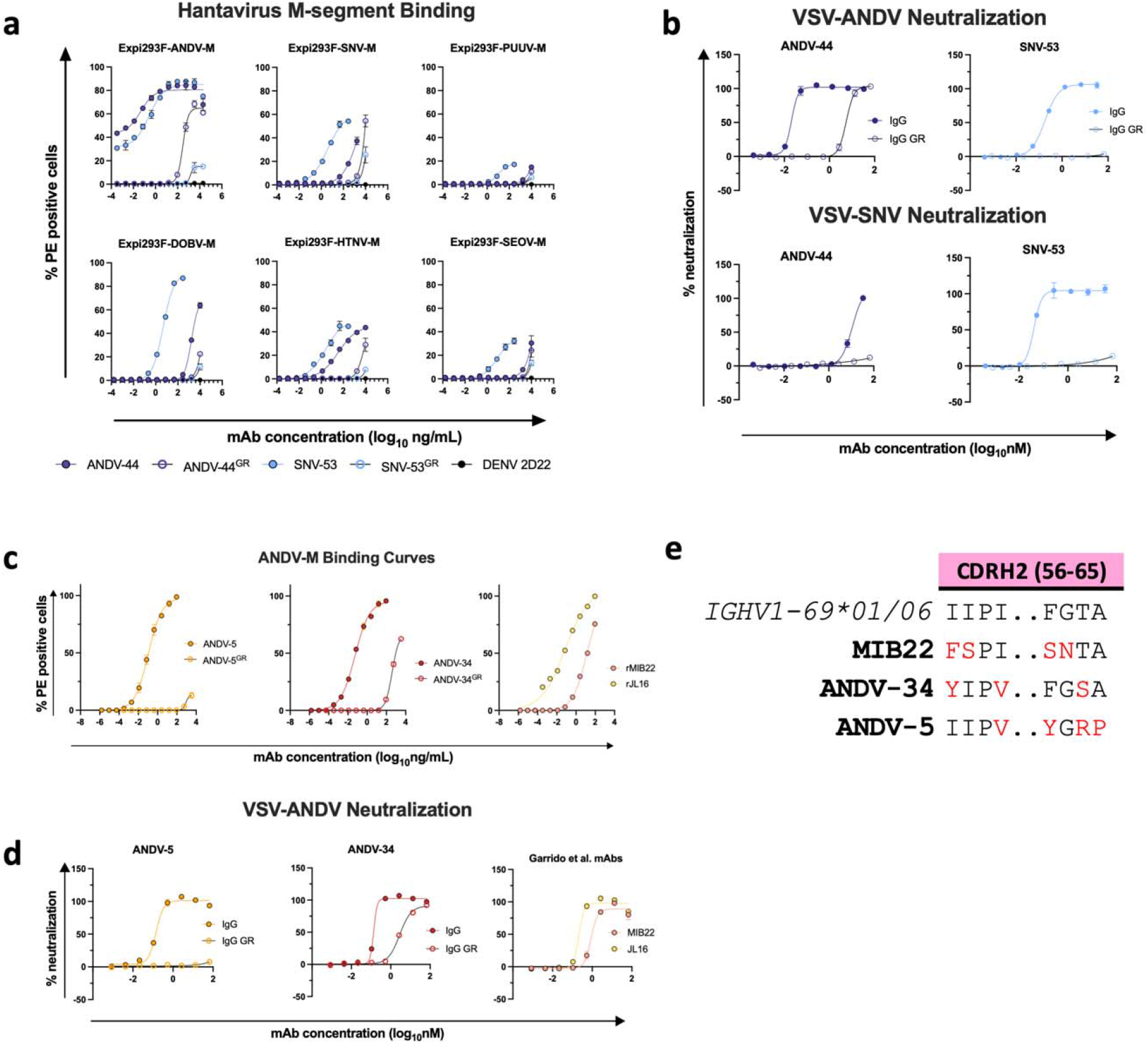
Reactivity and potency of germline revertant forms of NWH antibodies. **a.** Representative binding curves for all germline reverted forms of SNV-53 and ANDV-44 bnAbs to Expi293F cells transfected with ANDV, SNV, PUUV, DOBV, HTNV, or SEOV Gn/Gc. The value for % PE^+^ cells was determined by gating on cells stained only with secondary antibodies. Data shown are average values for technical replicates ± S.D. The experiment was performed 3 times independently with similar results; one experiment is shown. **b.** Representative neutralization curves for all bnAbs to VSVs bearing ANDV or SNV glycoproteins Gn/Gc. % Neutralization was measured using real-time cellular analysis and calculated by comparing CPE in the treatment wells with a cells only control well. Data shown are average values for technical replicates ± S.D. The experiment was performed 3 times independently with similar results. **c.** Representative binding curves for all ANDV-nAbs, including germline reverted forms, to Expi293F cells transfected with ANDV, as described in Extended Data Fig. 5a. **d.** Representative neutralization curves for all ANDV-nAbs to VSVs bearing ANDV, as described in Extended Data Fig. 5a. **e.** Alignment of MIB22, ANDV-34, and ANDV-5 CDHR2 sequences to the human germline IGHV1-69 gene. Somatically mutated residues are indicated in red. Alignment was generated using IMGT/DomainGapAlign.

**Extended Data Table 1.**
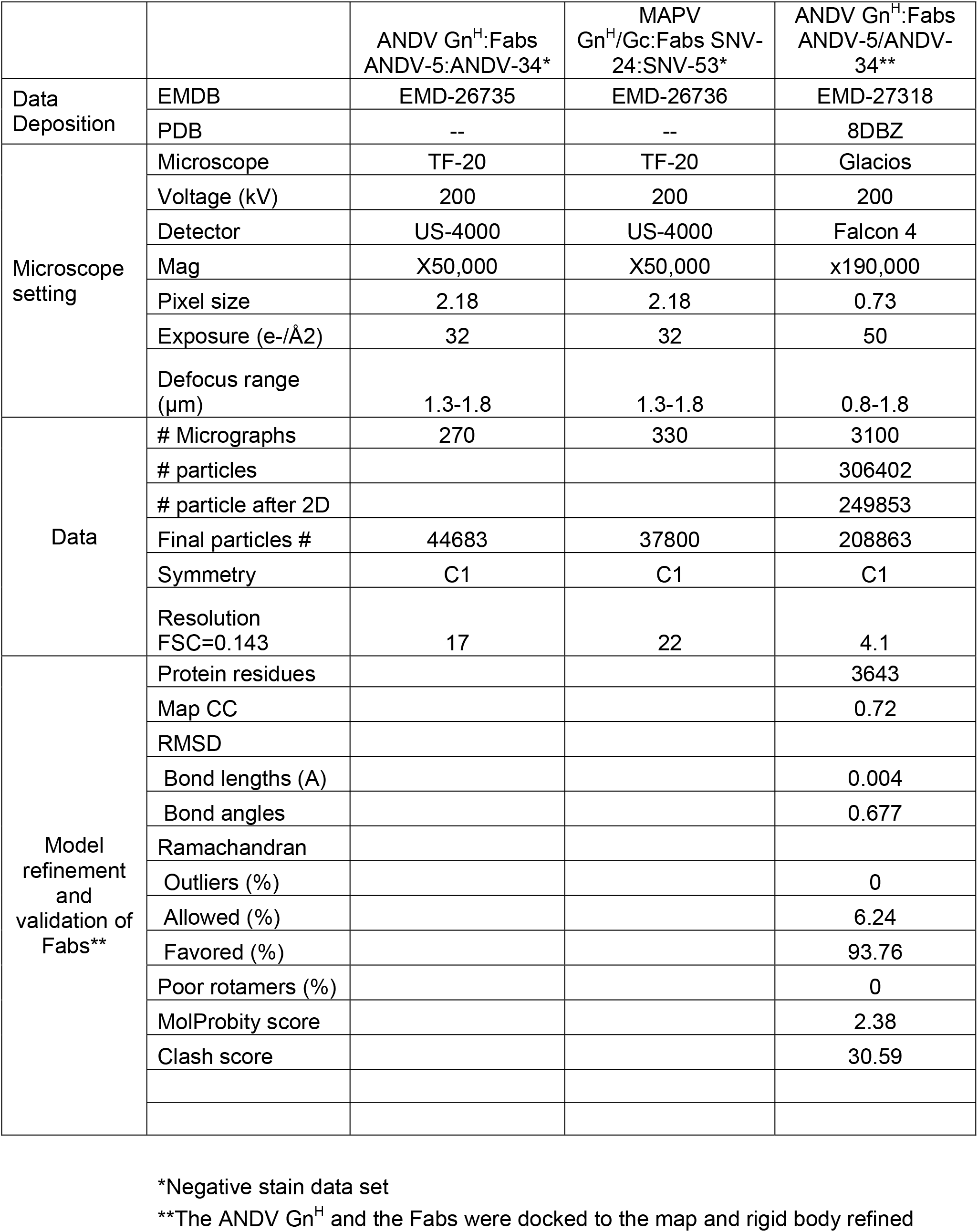
Summary table of electron microscopy statistics.

**Extended Data Fig. 6.**
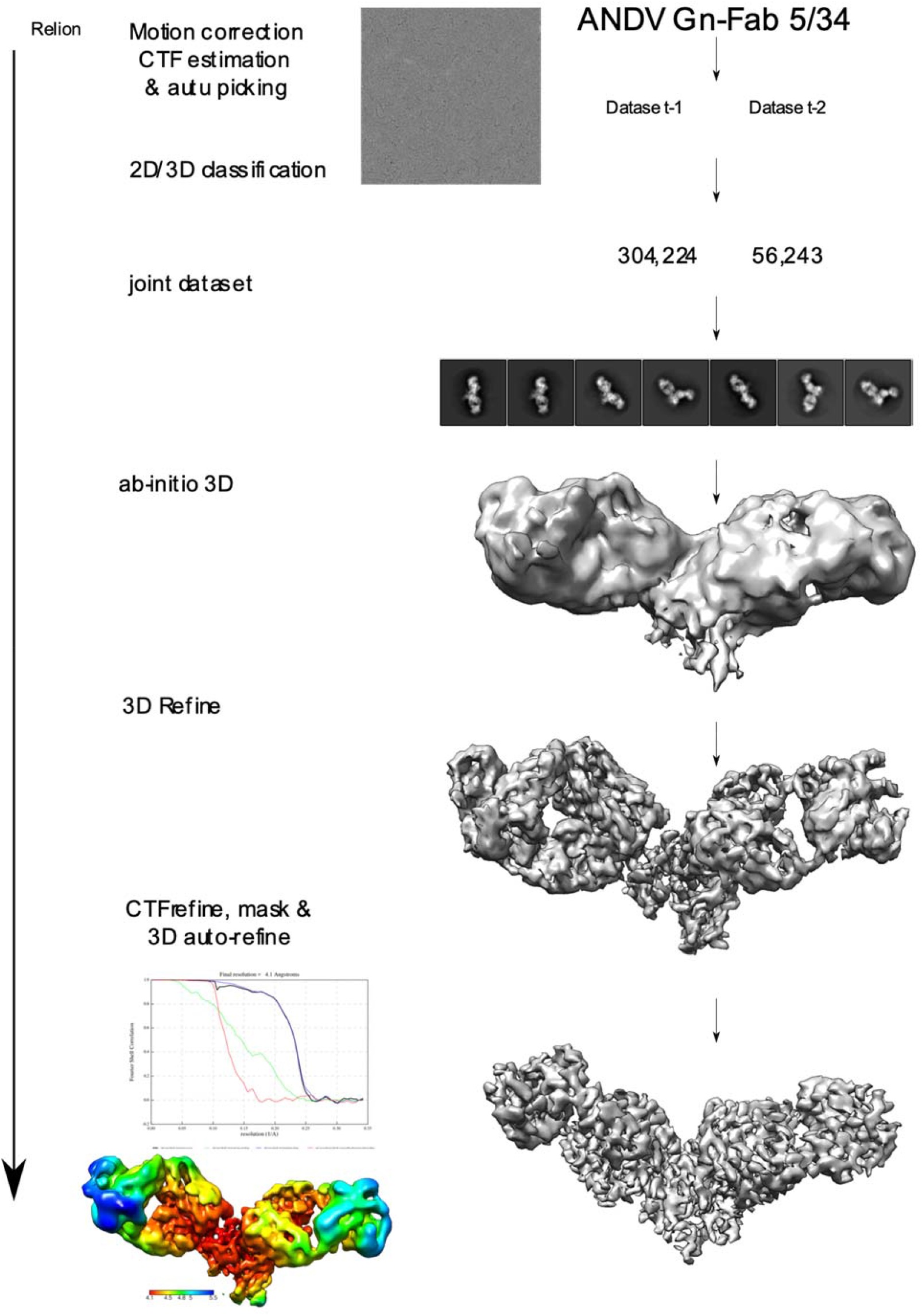
Workflow of cryo-electron microscopy processing for model of ANDV-5 and ANDV-34 Fabs in complex with ANDV Gn^H^.

**Extended Data Fig. 7.**
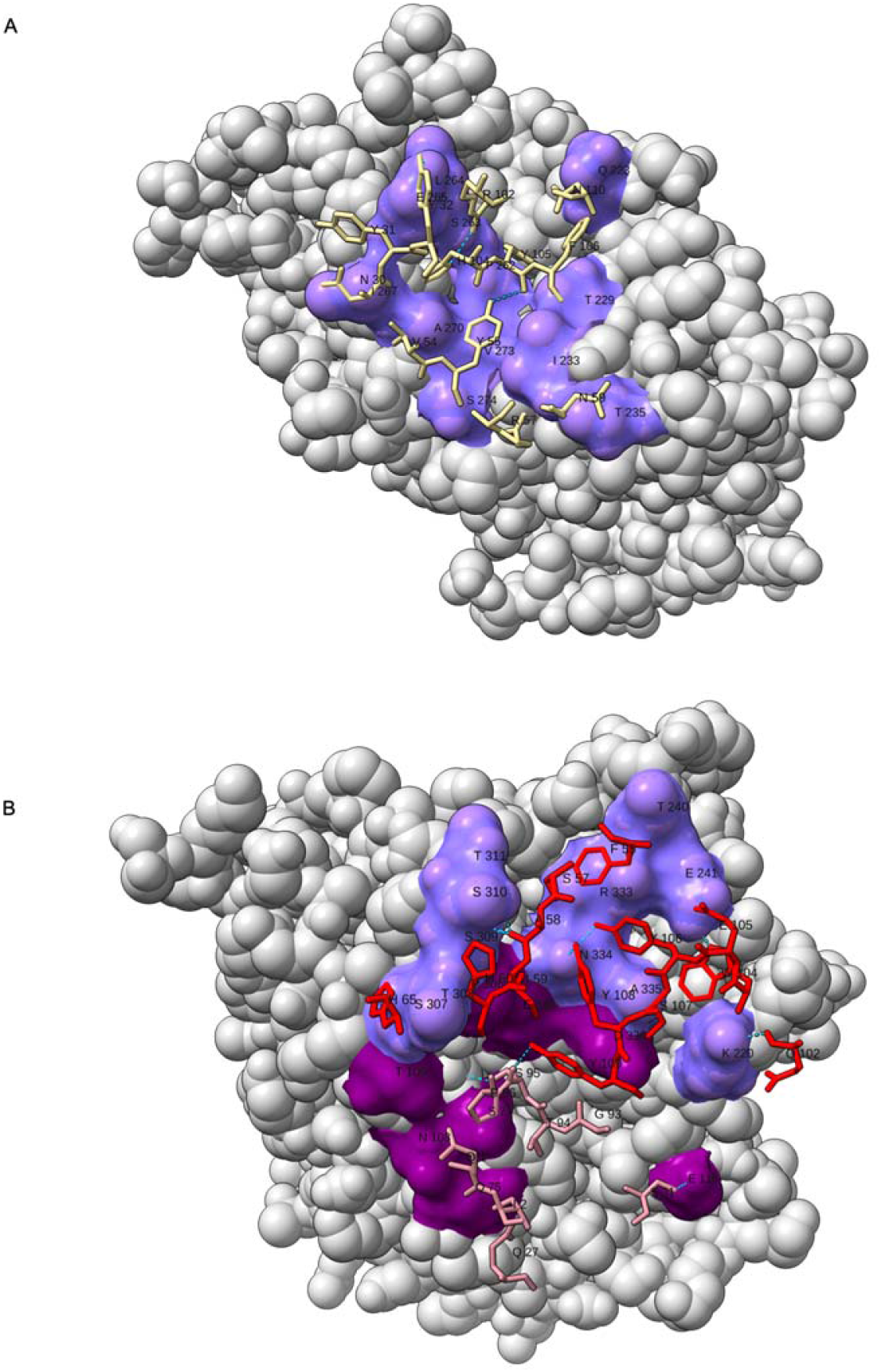
Residue interaction plot of Gn^H^ with ANDV-5 or ANDV-34. **a.** 3D representation of the interaction plot between Gn^H^ and ANDV-5. ANDV-5 is shown as sticks (yellow) and Gn as gray spheres with the contact residues in purple. Residues are labeled with a single letter and number. **b.** 3D representation of the interaction plot between Gn^H^ and ANDV-34. ANDV-34 is shown as sticks (HC-red and LC-pink) and Gn as gray spheres with the contact residues in purple (HC-light, LC-dark). Residues are labeled with a single letter and number.

**Extended Data Fig. 8.**
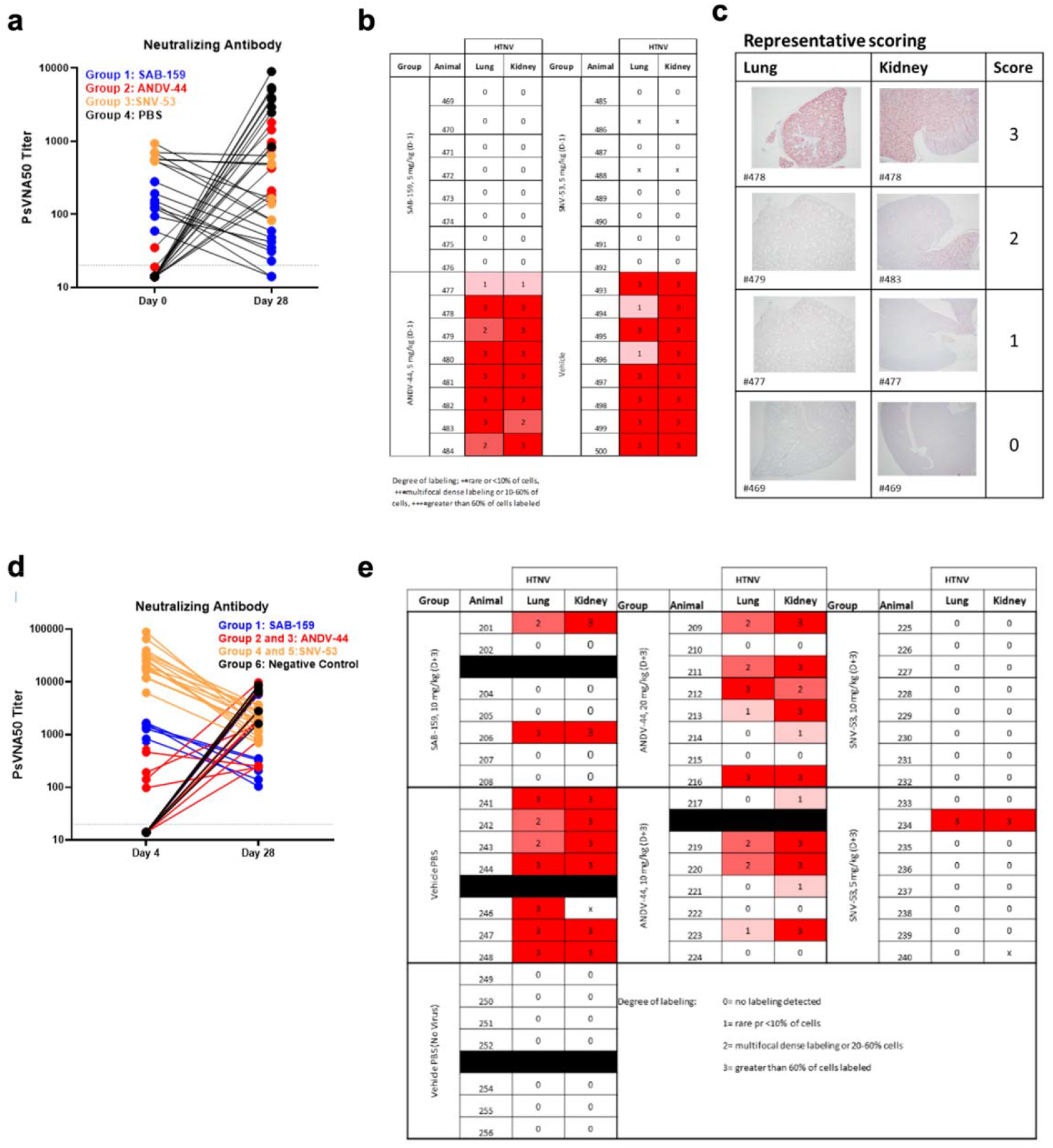
Pre- and post-exposure HTNV serum PsVNA50 data and pathology staining. **a.** Neutralizing antibody levels in serum for the four treatment groups at days 0 or 28 measured by PsVNA50 from the pre-exposure HTNV study. **b.** *in situ* hybridization (ISH) staining scores for individual animals from the pre-exposure HTNV study. **c.** Representative ISH staining in lung and kidney sections. Slides were scored by intensity ranging from 0 to 3, with median values presented. **d.** Neutralizing antibody levels in serum for the four treatment groups at days 4 or 28 measured by PsVNA50 from the post-exposure HTNV study. **e.** *in situ* hybridization (ISH) staining scores for individual animals from the post-exposure HTNV study.

